# Multifocal two-photon excitation fluorescence microscopy reveals hop diffusion of H-Ras membrane anchors in epidermal cells of zebrafish embryos

**DOI:** 10.1101/2022.10.25.513759

**Authors:** Radoslaw J. Gora, Redmar C. Vlieg, Sven Jonkers, John van Noort, Marcel J.M. Schaaf

## Abstract

Developments in fluorescence microscopy techniques have enabled imaging of individual fluorescently labelled proteins in biological systems, and in the current study, a single-molecule microscopy (SMM) technique has been applied *in vivo*, using the zebrafish embryo model. We have used multifocal two-photon excitation fluorescence microscopy (2PEFM) to study the dynamics of a GFP-fused H-Ras membrane-anchoring domain, GFP-C10H-Ras, in the epidermal cells of living embryos. In previous studies, a fast and a slow diffusing population of GFP-C10H-Ras molecules had been found. The application of the multifocal 2PEFM technique enabled us to focus on the slow diffusing population, which appears to occur in clusters that diffuse within microdomains of the epidermal cell membranes. Based on their mobility on a short timescale (≤ 1s) we could distinguish between a subpopulation that was diffusing and one that was virtually immobile. Owing to the multifocal 2PEFM imaging mode, we were able to dramatically reduce photobleaching which enabled us to follow the GFP-C10H-Ras particles over a prolonged time (> 3 s) and reconstruct their molecular trajectories of the diffusing subpopulation. These trajectories exhibited that the C10H-Ras particles continuously switch between a diffusing state and brief bursts of increased diffusion. As a result, they display an anomalous mobility pattern that can be referred to as hop diffusion. Taken together, this study demonstrates that multifocal 2PEFM offers a powerful approach to studying individual particles for prolonged periods of time, and that using this approach we were able to uncover the hopping behavior of GFP-C10H-Ras.

**SUMMARY STATEMENT:** By application of the two-photon excitation single-molecule microscopy to living zebrafish embryos, anomalous diffusion modes of individual H-Ras membrane anchors in epidermal cells were found.

## INTRODUCTION

Dissection of complex molecular processes, such as the dynamic behavior of proteins and lipids in the phospholipid membranes of living cells, requires an advanced technological approach that goes beyond traditional ensemble averaging techniques and a static molecular overview (Yokota, 2020). Single-molecule microscopy (SMM) techniques provide such advanced technology, and they have enabled scientists to detect individual molecules and quantitatively analyze their dynamics with unprecedented spatial and temporal resolution (e.g. Harms et al., 2001; Kusumi et al., 1993; Kusumi et al., 2012; Luo et al., 2020; Murakoshi et al., 2004). SMM is predominantly performed using advanced fluorescence microscopy approaches that include, among others, total internal reflection fluorescence (TIRF), highly-inclined and laminated optical sheet (HILO), and light-sheet fluorescence microscopy (LSFM) (e.g. Bernardello et al., 2021; Lommerse et al., 2006; Miller et al., 2018; Schaaf et al., 2009; Shashkova and Leake, 2017). Microscopy setups designed for imaging of individual fluorescent molecules are generally equipped with a laser beam used to excite fluorophores, and with a highly sensitive charged-couple device (CCD) or a complementary metal-oxide-semiconductor (CMOS) camera for efficient recording of the emitted photons (Axelrod, 2001). Numerous possibilities exist for the optimal fluorescent labelling of the molecules of interest, depending on the biological model and the properties of the imaging setup used (Chudakov et al., 2010; Seefeldt et al., 2008). Fluorescent labelling of proteins can, for example, be achieved through genetic fusion with traditional autofluorescent proteins, such as Green or Yellow Fluorescent Proteins (GFP, YFP), or photoactivable variants of these proteins (Li and Vaughan, 2018).

Until now, the majority of SMM studies have been performed in cultured eukaryotic, and often immortalized, cell lines (e.g. Chen et al., 2014; Gebhardt et al., 2013; Groeneweg et al., 2014; Ha et al., 1999; Keizer et al., 2019; Lommerse et al., 2004; Lommerse et al., 2005). These cell lines often show artefacts, have a limited translational value, and do not take into consideration the influence of cell-cell interactions as well as their interplay with the extracellular environment that occurs in an intact organism. Therefore, there is a need for the development of SMM techniques that can be applied to living organisms. For this purpose, we use zebrafish embryos, which are increasingly utilized in biomedical research as a vertebrate animal model, and are well suited for SMM imaging due to their optical clarity and because they are relatively easy to manipulate (Gore et al., 2018; Lu et al., 2015; Mione and Trede, 2010). The SMM techniques used for these *in vivo* studies should produce a limited background signal, make use of a large field of view (FoV), facilitate maintaining a biological sample alive over a prolonged period, and, preferably, not interfere with the natural biological processes of a sample by reducing, for instance, phototoxic and physicochemical stressors.

Thus far, we have extended SMM studies to living zebrafish embryos by investigating the mobility patterns of a yellow fluorescent protein (YFP) genetically fused to the human H-Ras protein, (YFP-H-Ras^WT^), its constitutively active and inactive genetic mutants (YFP-H-Ras^V12^ and YFP-H-Ras^N17^, respectively), and its membrane anchor (YFP-C10H-Ras) (Gora et al., 2021; Schaaf et al., 2009). The H-Ras protein is a member of the Ras protein family, which consists of small GTPases that activate intracellular signaling cascades, and thereby regulate crucial biological processes taking place in various cells, such as growth, proliferation, and differentiation (Malumbres and Barbacid, 2003). By use of the TIRFM technique and a dedicated mounting procedure, where the embryo tail tightly adheres to the coverslip, we succeeded in detecting individual, YFP-bound H-Ras proteins anchored to the apical cell membranes of cells in the outer epithelial layer of the epidermis, and in analyzing the dynamics of these molecules (Gora et al., 2021).

Using this approach, we were able to validate the results of previous studies, as we distinguished a fast- and a slow-diffusing fraction of H-Ras molecules, which both exhibited a confined type of diffusion. The fast-diffusing population contained most of the H-Ras proteins and was characterized by an initial diffusion coefficient that was approximately 10 to 15 times higher than the diffusion coefficient of the slow-diffusing H-Ras molecules. In addition, we discovered that a constitutively active H-Ras mutant (H-Ras^V12^) exhibited a significantly higher diffusion coefficient and a larger confinement area than wild type H-Ras. These increased dynamic parameters of the H-Ras^V12^ fast-diffusing fraction are generally considered to reflect preferential localization to specific membrane microdomains which have been suggested to result from local differences in lipid composition and the structure of the membrane cytoskeleton combined with transmembrane proteins anchored to the actin filaments (Fujiwara et al., 2002; Kusumi et al., 2012; Murakoshi et al., 2004). Therefore, we also studied the effect of zebrafish treatment with methyl-β-cyclodextrin (MBCD) and latrunculin B (LatB), which are known to disrupt the formation of a specific type of cholesterol-rich membrane microdomain, so-called lipid rafts, by cholesterol depletion or by inhibiting the formation of actin filaments respectively (Kwik et al., 2003). Both compounds significantly increased initial diffusion coefficients of the fast-diffusing fractions for the wild type H-Ras and its constitutively active, oncogenic mutant, H-Ras^V12^, together with their fraction sizes. The largest effects were observed for the non-activated wild type H-Ras, which is in line with its higher affinity for lipid rafts (Hancock and Parton, 2005; Prior et al., 2001).

Due to the limited penetration depth of the TIRFM technique, the applicability of this approach is restricted to proteins in the apical membrane of the outer epidermal cell layer. In the present study, we have explored the possibilities of a novel *in vivo* SMM approach using a multifocal two-photon excitation fluorescence microscopy (2PEFM) setup in order to develop a method with wider applicability (van den Broek et al., 2013). The 2PEFM is based on the near-simultaneous excitation of a fluorescent molecule with two photons, each containing approximately half the energy required to excite a molecule. The absorption of the two photons needs to occur within the fluorophore relaxation time of 10^−9^ s, which means that 2PEFM can only be performed using very high light intensities, which are achieved using pulsed laser sources with high peak intensities (in the range of kW-GW) and relatively low average laser power (2-5 W). This allows for lower laser beam absorption by a specimen being imaged and, therefore, reduced heating and phototoxic damage to the sample (Soeller and Cannell, 1999). Since the excitation light is generally in the near-infrared (NIR) wavelengths, and since biological tissues face less light scattering and absorption at NIR than at visible wavelengths, this makes 2PEFM a powerful technique for deep-tissue imaging. In addition, the background fluorescence is strongly reduced in 2PEFM, because the excitation of molecules is restricted to the confocal volume and the large anti-Stokes spectral difference between the excitation and emission light facilitates their spectral separation. To reduce the bandwidth limitations of the two-photon confocal microscopy, 2PEFM, we have combined it with multifocal scanning microscopy. In our multifocal microscope, a polygonal array of excitation light spots is generated in the image plane by passing the incident beam through a diffractive optical element (DOE). A two-dimensional fluorescence image is then created by scanning the array of spots across the sample plane within the camera exposure time, ensuring that the entire field of view (FoV) to be spanned by such an array of spots is homogenously illuminated.

In the present study, we have used a multifocal 2PEFM setup to analyze the mobility pattern of GFP fused to the human H-Ras membrane-anchoring domain (referred to as GFP-C10H-Ras) in epidermal cells of two-day-old zebrafish embryos. The multifocal 2PEFM technique enabled us to visualize GFP-C10H-Ras molecules of the slow-diffusing fraction. We show that these molecules occur in clusters that move through the plasma membrane, and that this population consists of a diffusing and an immobile fraction. By following the mobility of the diffusing molecules for at least 3 seconds (i.e., 15 frames), we showed that these diffusing GFP-C10H-Ras molecules alternate between a diffusing state and a state with an increased diffusion rate which is referred to as ‘hopping’.

## RESULTS

### Imaging of GFP-C10H-Ras molecules in epidermal cells of zebrafish embryos using 2PEFM

To analyze the *in vivo* mobility pattern of the H-Ras membrane anchor, the GFP fusion protein of the H-Ras membrane-anchoring domain, GFP-C10H-Ras, was studied in 2 days-post-fertilization (dpf) zebrafish embryos using a 2PEFM setup. We focused our observations on the apical membrane of the outer epidermal cell layer, i.e., the superficial stratum, in the tail fin of the embryos from a *Tg(bactin: GFP-C10H-Ras)*^*vu119*^ transgenic line (Fig. 1). The embryos from this line stably express the GFP-tagged H-Ras membrane anchor in virtually all cells, including the epidermal cells in their tail fin, which is illustrated by a homogenous GFP-C10HRas localization in pentagonal and hexagonal cells of the outer epidermal cell layer (Fig. 1). The 2 dpf zebrafish embryos were placed on a coverslip and the tail fin region was covered with a 0.75-mm-thick sheet of agarose, which was used to gently press the tail fin towards the surface and, therefore, further reduce the possible movement of the anaesthetized embryo on the coverslip during the imaging. The rest of the zebrafish body was covered with a drop of water. Zebrafish vital functions, including heartbeat and the blood flow in its cardiovascular system, were controlled under a stereofluorescence microscope post-imaging.

**FIGURE 1:**
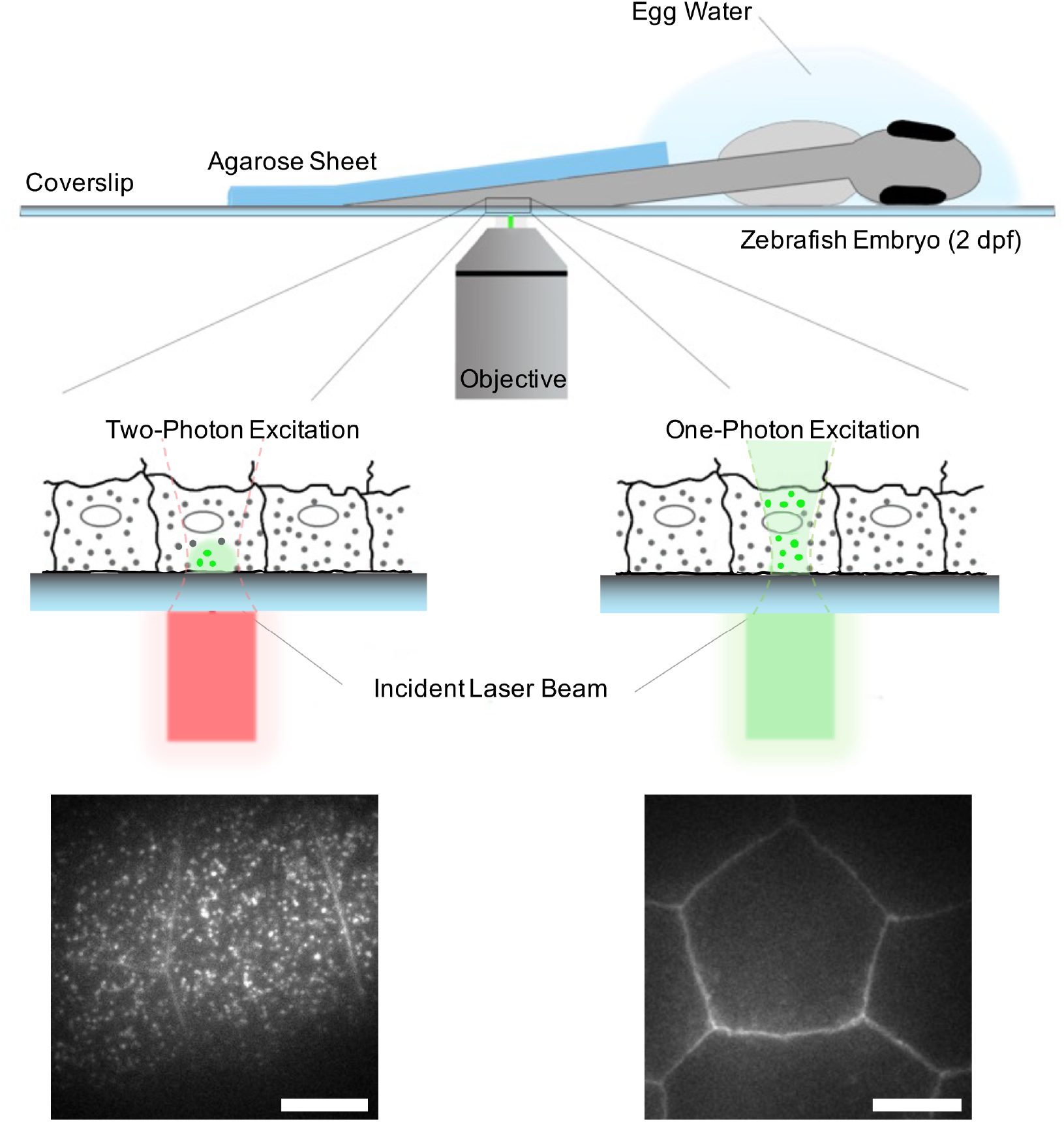
Schematic overview of SMM applied to a living zebrafish embryo. A zebrafish embryo from a Tg*(bactin: GFP-C10H-Ras)*^*vu119*^ transgenic line stably expressed the GFP-tagged H-Ras anchor in the zebrafish tailfin. At 2 dpf, it was placed on a coverslip coated with poly-L-lysine and coated with a drop of egg water. The tail region of the embryo was covered with a 0.75-mm-thick agarose (2%) sheet. On the lower right part of the figure, a picture of the outer layer of the epidermis is presented, showing the fluorescent signal of GFP-C10H-Ras in the cell membranes. This picture was taken using a one-photon excitation fluorescence microscopy technique. Morphologically, the cells in this layer are homogenous and are characterized by pentagonal and hexagonal shapes. On the lower left part, a 2PEFM image of an area with the same FOV is presented, with clearly visible localizations of the GFP particles within the zebrafish outer epidermal layer. Scale bar: 5 μm.

Image sequences with a time lag of 200 ms were acquired using the 2PEFM setup. As a first step in the analysis, we investigated the characteristics of the fluorescence intensity peaks detected in our images (Fig. 2A). We fitted the intensity peaks to a Gaussian surface, using custom-made software developed previously (Harms et al., 2001; Schütz et al., 1997a). Interestingly, we observed particles with varying maximum intensity counts, as shown in the density plots for these parameters (Fig. 2BCD), indicating that GFP particles of varying molecular compositions occur in our molecular populations which indicates clustering of GFP-C10H-Ras molecules. The extremum of the FWHM density plot equaled 327 nm, whereas the extremum of the density plot for the maximum intensity value equaled 893 counts.

**Figure 2:**
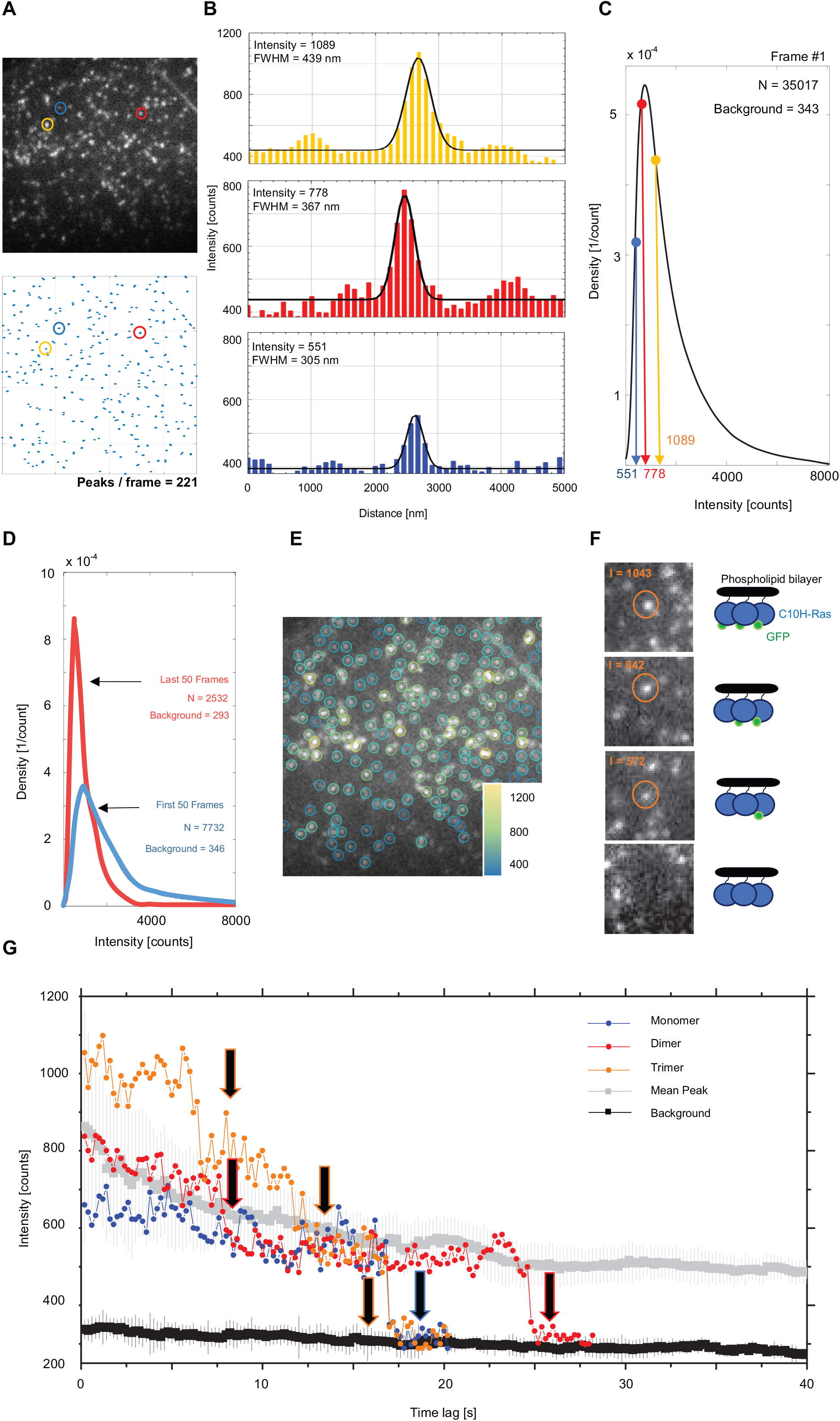
Structural analysis of the H-Ras anchor populations observed in the 2PEFM setup. **(A)** 2PEFM image of the GFP particles with a selection of three particles with differing intensities in circles (upper panel); localization map of the GFP particles in the same image, acquired through a custom-made MATLAB software (lower panel). **(B)** 2D Gaussian curves fitted onto the three particles selected in (A). The graph in yellow denotes an intensity profile example of a trimeric GFP, the one in red a dimeric GFP, and the one in blue a single GFP molecule. **(C)** Examples of density plots of the intensity values in the first frame of a time lapse. Values of the three particles selected in (A) are overlayed, indicating the wide spectrum of GFP multiplicities present in our molecular population. **(D)** Density plots of the intensity counts in the first and the last 50 frames of the time lapse consisting of 600 individual frames. Notable is the switch in the most abundant GFP populations, from polymeric GFPs in the first 50 frames to monomeric GFP molecules. **(E)** The 2PEFM image from (A) with a GFP particles attributed a circular selection of 5 pixels in diameter, with intensity of each selection determined by the pixel with the highest intensity count within the selection, performed by an ImageJ plug-in, TrackMate (lower panel). **(F)** Visualization of a photobleaching event, with an example of a GFP trimer. Three distinctive intensity levels are depicted, denoting a step-like decrease of their intensity count. **(G)** Single-step photobleaching experiment. Intensity values of the GFP circular selections were acquired from five individual 2PEFM time lapses, and are depicted as the means with the corresponding standard error values (in grey). The background intensity is defined as a mean intensity count of all the pixels falling outside the peak selections within each frame, and are depicted as the means with the corresponding standard error values (in black). Molecules selected in (A) are overlayed and followed over time, indicating their step-like intensity drop. I, intensity count; N, number of localized molecules; Background, mean background intensity.

In order to study possible changes in the peak characteristics over time, we generated the density plots for the maximum intensities of the peaks in the first and the last 50 frames of five different image sequences, which all comprised a total of 600 frames. A clear difference in the density plot of the maximum intensity values was observed, as the extremum of the density dropped from 875 counts (median of 943) in the first 50 frames to 567 (median of 582) in the last 50 frames of the time lapses (Fig. 2D). There was a wide distribution in the peak intensity in the first frames of the image sequence as a result of the clustering of molecules, but these intensities converge to a level of around 500-600 counts in the final frames, most likely as a result of photobleaching of molecules within the clusters which near the end of the image sequence mainly contain only a single fluorescent molecule.

In a subsequent, more detailed analysis of clustering and photobleaching events, all GFP peaks were identified using the ImageJ plug-in TrackMate (Tinevez et al., 2017). In the plug-in, a circular selection with a diameter of 5 pixels was drawn around each of the located GFP particles. The located GFP particles were sorted according to their intensity levels, defined by the pixel with the highest intensity value within the circular selection (Fig. 2E). Using this analysis, we studied the alterations in the characteristics of the individual intensity peaks over time (Fig. 2F). The circular selections representing GFP clusters were followed over time, and their maximum intensity counts were averaged.

The background intensity of an image was defined as the average intensity count of all pixels that fell outside the circular selections. Then, we followed three peaks with varying initial intensity values over time, and it appeared that all three peaks exhibited step-wise photobleaching. The intensity level prior to the final photobleaching step was in the range of 500-600 intensity counts (Fig. 2G), indicating that this intensity level corresponded to a population of single fluorescent GFP molecules. This finding was further confirmed by the analysis of the average peak intensity count per image, which decreased over time and stabilized during the final stage of the time-lapse around this intensity range (Fig. 2G, grey line). The outcome was in line with the single GFP signals being the most abundant among all detected intensity peaks at this stage, as a result of the photobleaching. Based on these results, we concluded that it is possible to detect single GFP molecules with our 2PEFM setup, and that most of the detected GFP-C10H-Ras molecules formed clusters in the plasma membrane. Subsequently, the mean time until photobleaching was determined for the GFP molecules imaged with either TIRF or 2PEF microscopy techniques, by plotting the number of photobleaching events as a function of time (Fig. 3AB). Fitting of the mean lifetime for particle decay function, 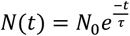, with *N*_0_ as an initial quantity and *t* as time, provided mean lifetimes *τ* of 0.067 s for TIRF and 0.783 s for 2PEF (2.7 and 3.9 times the integration time respectively). These data show that 2PEF microscopy results in significantly delayed photobleaching compared to TIRF.

**FIGURE 3:**
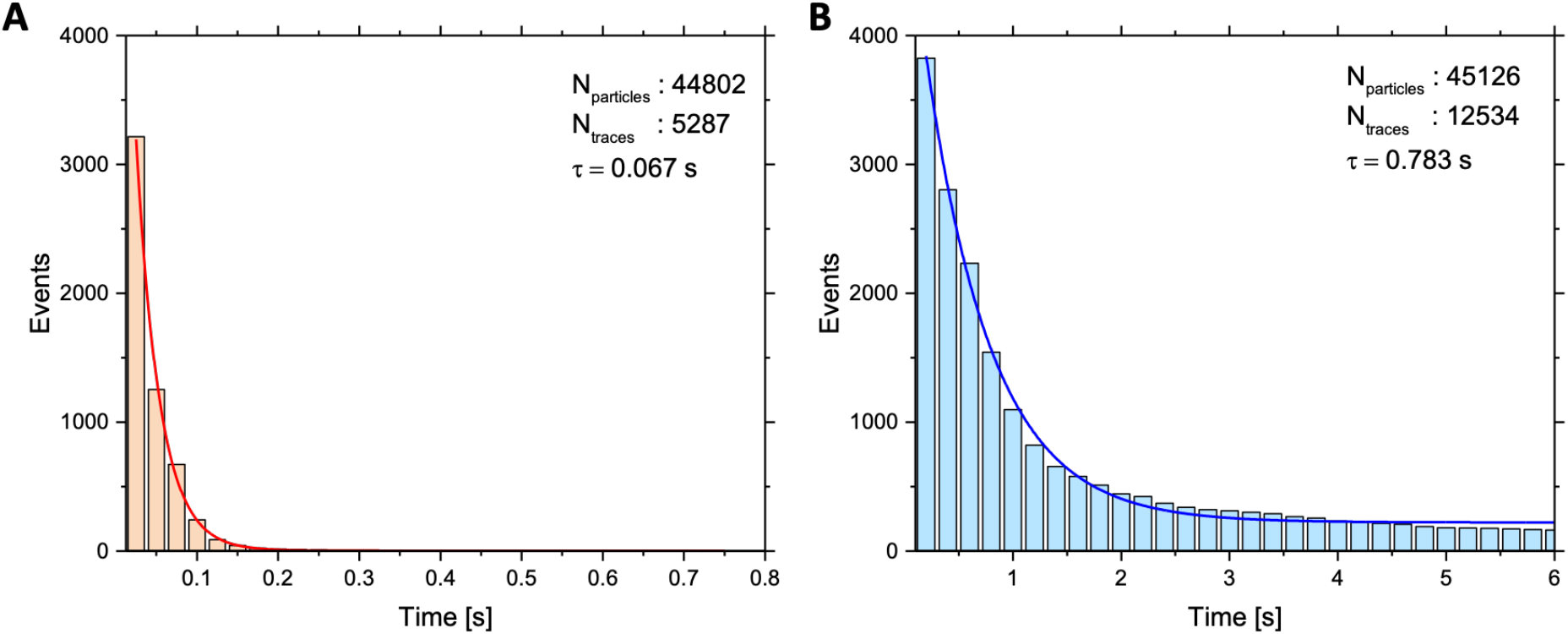
Time until photobleaching for the fluorescent molecules imaged with (A) TIRFM and (B) 2PEFM. In the graphs, the number of photobleaching events as a function of time are presented. A one-exponential decay function is overlayed to establish an average lifetime *τ* of fluorophores using the two microscopy techniques **(A)** The average lifetime *τ* of fluorophores imaged with TIRFM with an integration time of 0.025 s equaled 0.067 s. **(B)** The average lifetime *τ* of fluorophores imaged with 2PEFM with an integration time of 0.200 s equaled 0.783 s.

### Analysis of the mobility pattern of GFP-C10H-Ras molecules in epidermal cells of zebrafish embryos using 2PEFM

The mobility patterns of the GFP-fused H-Ras membrane anchoring domains were analyzed using Particle Image Correlation Spectroscopy (PICS) software (Semrau and Schmidt, 2007). Further, using the PICS analysis, correlations between the locations of molecules in consecutive frame pairs were determined. Based on this analysis, cumulative probability distributions of the displacements were generated for each time lag. These curves were then fitted to a one-, (Eq. 1) a two-(Eq. 2), or a three-population (Eq. 3) model (Fig. 4A; for further reference, see Materials and Methods section). The one-population model fitted the 2PEFM data with a Pearson’s correlation coefficient of 0.948, whereas the two-population model much better reflected the mobility patterns of the GFP-C10H-Ras, and the correlation coefficient of the fit equaled 0.999. Since the three-population models yielded a similar correlation coefficient, we used the two-population model for further analysis, which best fitted the experimental data with the least number of populations. Using this model, which suggests the presence of two fractions of molecules with different diffusion rates, the relative size of the fast-diffusing fraction (*α*) and the mean squared displacements of the fast- and slow-diffusing fractions (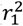 and 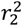, respectively) were determined. By using a multistep analysis to investigate the GFP-C10H-Ras dynamics, these parameters were determined at five different time lags: 200, 400, 600, 800, and 1000 ms.

Subsequently, the parameters 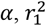, and 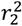 were plotted against the time lag (Fig. 4BCD). The relative size of the GFP-C10H-Ras fast-diffusing fraction *α* was stable over all time lags (Fig. 4B, Table 1), and equaled 49.5 ± 0.4%. For both the fast- and the slow-diffusing fraction, the plots presenting the mean squared displacements (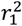 and 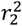)as a function of the time lag reached a plateau (Fig. 4CD). Hence, we fitted these curves to a confined diffusion model (Eq. 4), in which the H-Ras membrane anchors move with an initial diffusion coefficient *D*_0_ and their mobility is confined to a squared area with sides of length *L* (Bobroff, 1986; Lingwood and Simons, 2010; Schaaf et al., 2009). Based on our analysis, the *D*_0_ of the fast-diffusing fraction equaled 0.066 ± 0.005 μm^2^ s^-1^ and the size of its confinement area *L* equaled 253 ± 3 nm. The slow-diffusing fraction of the GFP-C10H-Ras hardly diffused with a *D*_0_ of 0.80 × 10^−3^ ± 0.10 × 10^−3^ μm^2^ s^-1^, and was confined to a smaller area (*L* of 94 ± 5 nm). Since the mean squared displacements of this population were similar to the offset value in this plot (0.0031 μm^2^, based on the positional accuracy of ∼28 nm), the molecules in this population may be considered immobile. When the data were analyzed using the one-population model, the *D*_0_ of this single population equaled 0.116 ± 0.011 μm^2^ s^-1^ and the confinement area size *L* equaled 302 ± 54 nm (Fig. S1).

**TABLE 1:** Summary of the GFP-C10H-Ras mobility patterns acquired using the PICS analytical method.

| Experiment | $D_{0FD}$ [$\mu\text{m}^2 \text{s}^{-1}$] | $D_{0SD}$ [$\mu\text{m}^2 \text{s}^{-1}$] | $L_{FD}$ [nm] | $L_{SD}$ [nm] | $\alpha$ [%] |
| --- | --- | --- | --- | --- | --- |
| **1.1. Mobility patterns of GFP-C10H-Ras in zebrafish embryos: 2PEFM imaging** |  |  |  |  |  |
| Zebrafish embryo | $0.066 \pm 0.005$ | $(0.080 \pm 0.010) \cdot 10^{-2}$ | $253 \pm 3$ | $94 \pm 5$ | $49.5 \pm 0.4$ |
| **1.2. Mobility patterns of GFP-C10H-Ras in zebrafish embryos: comparisons with TIRFM imaging** |  |  |  |  |  |
| Time lag: 25 ms | $1.46 \pm 0.16$ | $0.152 \pm 0.047$ | $468 \pm 4$ | $103 \pm 5$ | $66.0 \pm 5.3$ |
| Time lag: 200 ms | $0.22 \pm 0.03$ | $(0.312 \pm 0.101) \cdot 10^{-2}$ | $553 \pm 2$ | $128 \pm 2$ | $52.5 \pm 1.1$ |
| Time lag: 25 ms: Three-population model | $1.15 \pm 0.03$ | $0.306 \pm 0.045$ ($D_{0SD}$)<br>$0.042 \pm 0.007$ ($D_{0IM}$) | $581 \pm 8$ | $286 \pm 16$ ($L_{SD}$)<br>$102 \pm 18$ ($L_{IM}$) | $51.0 \pm 3.2$ ($\alpha_1$)<br>$39.2 \pm 2.3$ ($\alpha_2$) |
| **1.3. Mobility patterns of GFP-C10H-Ras in zebrafish embryos: impact of TIRFM differing excitation power** |  |  |  |  |  |
| 50% of the total power | $0.91 \pm 0.14$ | $0.092 \pm 0.010$ | $444 \pm 3$ | $108 \pm 47$ | $54.8 \pm 6.3$ |
| 30% of the total power | $1.22 \pm 0.22$ | $0.089 \pm 0.008$ | $475 \pm 5$ | $112 \pm 27$ | $60.7 \pm 9.8$ |
| 10% of the total power | $0.76 \pm 0.02$ | $0.043 \pm 0.010$ | $536 \pm 3$ | $184 \pm 13$ | $61.7 \pm 15.7$ |
| **1.4. Mobility patterns of GFP-C10H-Ras in zebrafish embryos: effect of vehicle, LatB and MBCD treatments** |  |  |  |  |  |
| GFP-C10H-Ras in vehicle | $0.051 \pm 0.006$ | $(0.080 \pm 0.003) \cdot 10^{-2}$ | $237 \pm 3$ | NA | $49.9 \pm 0.3$ |
| GFP-C10H-Ras + LatB | $0.039 \pm 0.007$ | $(0.055 \pm 0.006) \cdot 10^{-2}$ | $213 \pm 5$ | NA | $53.7 \pm 0.9$ |
| GFP-C10H-Ras + MBCD | $0.110 \pm 0.020$ | $(0.085 \pm 0.005) \cdot 10^{-2}$ | $277 \pm 2$ | NA | $55.5 \pm 0.2$ |

$D_{0FD}$ , $D_{0SD}$ , $D_{0IM}$ – Initial diffusion coefficients for fast-, slow-diffusing and immobile fractions, respectively; $L_{FD}$ , $L_{SD}$ , $L_{IM}$ – Sizes of the confinement area of fast-, slow-diffusing, and immobile fractions, respectively; $\alpha$ – fast-diffusing fraction size.

**TABLE 2:** Statistical analysis performed for the values of mean squared displacements and slow-diffusing fraction sizes obtained experimentally for the time lag of 200 ms.

| Statistical significance<sup>1</sup> | Mean squared displacement: slow-diffusing fraction ($r_1^2$) | Mean squared displacement: immobile fraction ($r_2^2$) | Fraction size: slow-diffusing fraction ($\alpha$) |
| --- | --- | --- | --- |
| GFP-C10H-Ras in zebrafish embryos treated with LatB and MBCD | P = 0.091 | P = 0.362 | P = 0.324 |
| GFP-C10H-Ras in zebrafish embryos imaged with 2PEFM and TIRFM setups<sup>2</sup> | P(2PEFM, TIRFM<sup>25</sup>) < 0.001<br>P(2PEFM, TIRFM<sup>200</sup>) < 0.001<br>P(TIRFM<sup>25</sup>, TIRFM<sup>200</sup>) < 0.001 | P(2PEFM, TIRFM<sup>25</sup>) < 0.001<br>P(2PEFM, TIRFM<sup>200</sup>) = 0.619<br>P(TIRFM<sup>25</sup>, TIRFM<sup>200</sup>) = 0.005 | P(2PEFM, TIRFM<sup>25</sup>) < 0.001<br>P(2PEFM, TIRFM<sup>200</sup>) = 0.164<br>P(TIRFM<sup>25</sup>, TIRFM<sup>200</sup>) < 0.001 |
| GFP-C10H-Ras in zebrafish embryos imaged with TIRFM at differing excitation powers | P > 0.132 | P > 0.262 | P = 0.294 |

<sup>1</sup>For comparisons of protein mobility patterns in zebrafish embryos between different chemical treatments, microscopy settings, and TIRFM laser excitation powers, a one-way ANOVA was used. In case the differences between groups were significant in the one-way ANOVA, results of multiple groups comparison are shown, performed by a Tukey's range posthoc test.
<sup>2</sup>In the case of TIRFM imaging with the time lag of 25 ms, values of $\alpha$ , $r_1^2$ , and $r_2^2$ for statistical analysis were obtained experimentally for the time lag of 25 ms.
NS – non-significant (P > 0.05); \* – P < 0.05; \*\*\*\* – P < 0.001.

**FIGURE 4:**
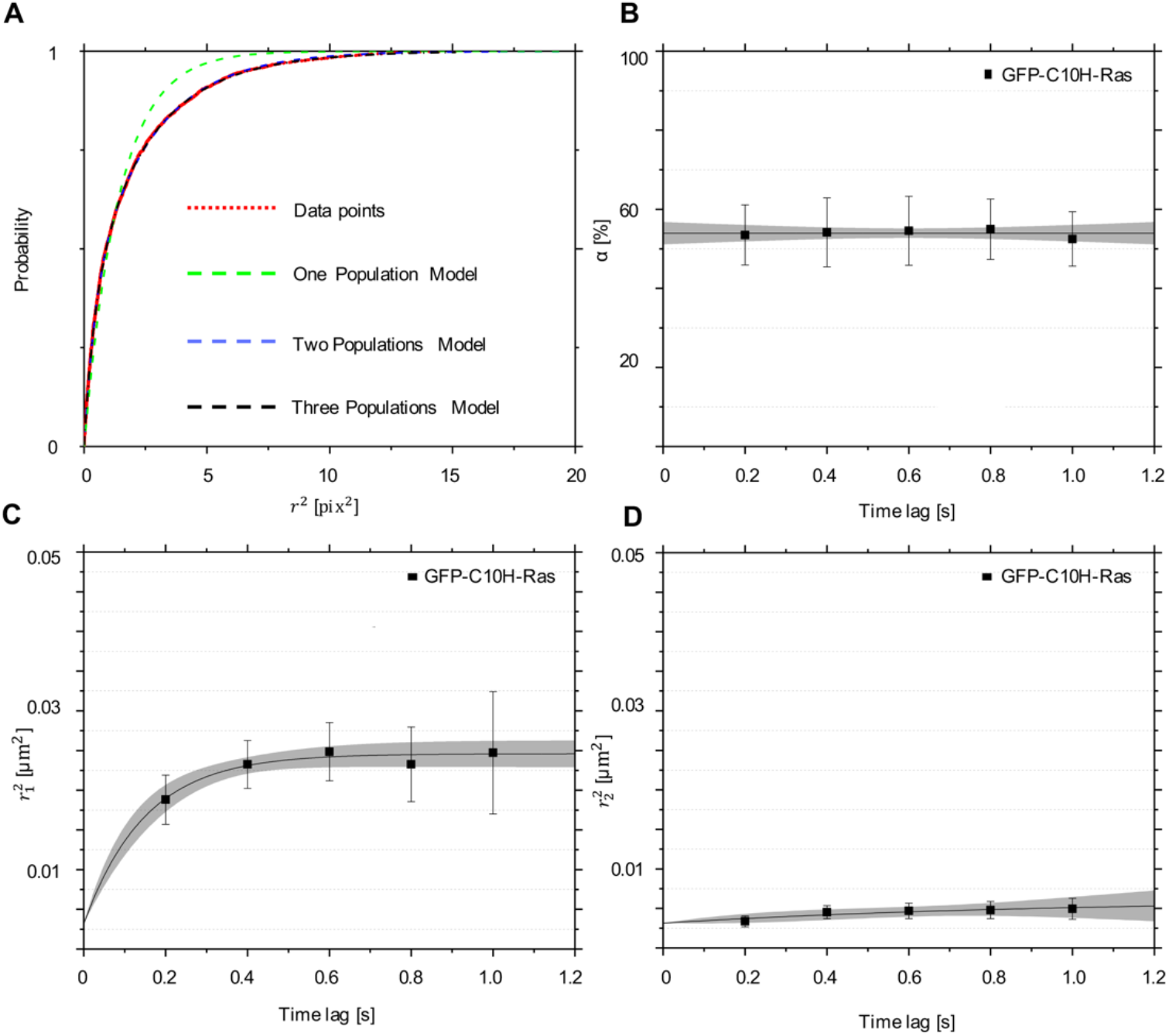
Mobility patterns of GFP-C10H-Ras in epidermal cells of the zebrafish embryos. **(A)** Cumulative probability distribution function of the GFP-C10H-Ras squared displacements calculated for the 2PEFM datasets. Data points are shown in red, and their population fits in dashed green for a one-population model, in dashed blue for a two-population model, and in dashed black for a three-population model. No significant difference was observed between the fitting of the two- and the three-population models, which both best fitted the cumulative probability distribution. Fitting of the data points to the two-population model allowed for calculation of the mean squared displacements of these populations (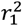 and 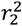). This procedure was repeated for each of the time lags used. **(B, C, D)** Plots representing mobility patterns of the GFP-C10H-Ras inside the zebrafish embryos from the Tg(bactin: GFP-C10H-Ras)^vu119^ transgenic line. (**B**) Fraction size of the fast-diffusing population (*α*), plotted against the time lag. **(C)** Mean squared displacements plotted against the time lag for the fast-diffusing fraction (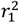) **(D)** Mean squared displacements plotted against the time lag for the slow-diffusing fraction (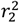). To establish the values of dynamic parameters for the GFP-C10H-Ras from the Tg(bactin: GFP-C10H-Ras)^vu119^ line, 5 different embryos were imaged on each of the 5 different experimental days. Parameters obtained upon curve fitting are presented in Table 1. Each data point is presented in the form of a mean ± s.e.m., and the 95% c.i. of the mathematical fit is shown. Shapiro-Wilk statistical test was performed to check for the normality of the data set. Statistical analysis for all treatment groups was performed using a one-way ANOVA (P-value 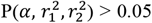 at a *t*_*lag*_ of 200 ms) with a Tukey’s range posthoc test (for details of the post-hoc test results, see Table 2).

Interestingly, the initial diffusion coefficients and confinement area sizes of the two GFP-C10H-Ras subpopulations that were determined in the zebrafish epidermal cells using 2PEFM imaging did not correspond to the findings previously obtained for YFP-C10H-Ras in the same cells using a TIRFM setup (Gora et al., 2021; Schaaf et al., 2009). In particular, the fast-diffusing fraction, as identified in the previous TIRFM experiments, was not detected in the current study. In addition, the slow-diffusing YFP-C10H-Ras population previously identified using TIRFM showed a mobility pattern highly similar to that of the fast-diffusing subpopulation that was discerned using 2PEFM imaging in this study (Gora et al., 2021; Schaaf et al., 2009). This suggests that using the current approach we are not able to detect the fast-diffusing population of GFP-C10H-Ras molecules that we previously observed using TIRFM. Moreover, the slow-diffusing population that was previously identified in our TIRFM experiments was observed in the present study, and this population could be further divided into a slow-diffusing and an immobile subpopulation based on the novel data.

### Comparison of the GFP-C10H-Ras dynamics between 2PEFM and TIRFM with different temporal resolutions

In order to determine whether the absence of the fast subpopulation of GFP-C10H-Ras molecules is a result of the temporal resolution of the 2PEFM, we determined dynamic parameters for GFP-C10H-Ras by using the TIRFM setup and imaging the same batches of zebrafish embryos. We imaged embryos with a time lag of 25 ms, which had also been used in previous studies (Gora et al., 2021; Schaaf et al., 2009), and a time lag of 200 ms, which corresponds to the time lag used in the 2PEFM experiments (Fig. 5A). With both time lags, we could distinguish a fast- and a slow-diffusing fraction of molecules using the analysis based on a two-population model (Fig. 5B), and all fractions showed confined diffusion, but generally, the mobility of GFP-C10H-Ras decreased with increasing time lag (Fig. 5CD). The size of the fast-diffusing fraction decreased significantly (66.0 ± 5.3% at 25 ms, and 52.5 ± 1.1% at 200 ms), as did the initial diffusion coefficients *D*_0_ for the fast-diffusing fraction (1.46 ± 0.16 μm^2^ s^-1^ at 25 ms, and 0.22 ± 0.03 μm^2^ s^-1^ at 200 ms). On the other hand, the size of the confinement area *L* for GFP-C10H-Ras increased (468 ± 4 nm at 25 ms and 553 ± 2 nm at 200 ms). For the slow-diffusing fraction, the initial diffusion coefficients decreased with increasing time lag (0.152 ± 0.047 μm^2^ s^-1^ at 25 ms and 0.312·10^−2^ ± 0.101·10^−2^ μm^2^ s^-1^, whereas the size of the confinement area increased (103 ± 5 nm at 25 ms and 128 ± 2 nm at 200 ms). Taken together, these findings demonstrate that the temporal resolution of the microscopy setup affects the dynamic parameters determined for GFP-C10H-Ras. In particular, an increase in the time lag results in lower mobility. It is most likely that due to a longer illumination time, the signal of the fast-diffusing molecules is spread over a larger area, generating a motion blur, and making it impossible to distinguish single fluorescent signal spots above the background signal.

Thus, we demonstrated that the time lag of 200 ms was largely responsible for the decreased mobility that we determined in our 2PEFM study compared to our previous TIRFM studies. When we compared the results from the TIRFM and 2PEFM experiments, both using a 200 ms time lag, we found no significant differences between the population sizes, and the mobility pattern of the slow-diffusing populations (Table 1, 2). However, the fast-diffusing population observed in the 2PEFM experiment still showed a lower diffusion coefficient and a smaller confinement area than its equivalent observed by TIRFM, so the lower mobility observed of this fraction using 2PEFM cannot entirely be explained by the lower time resolution.

**FIGURE 5:**
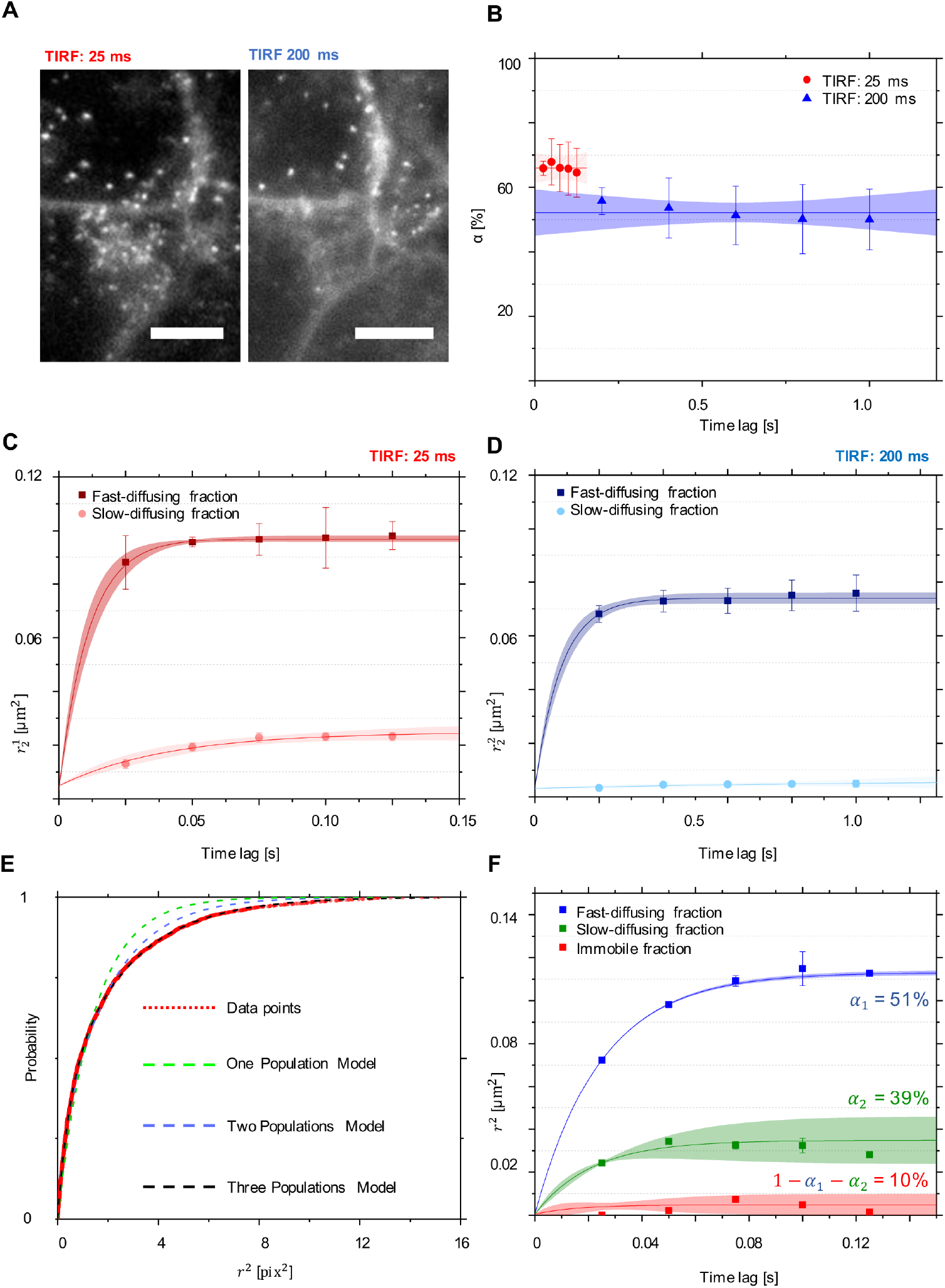
GFP-C10H-Ras mobility patterns: differences between time lags in the TIRFM setup. Results of different multi-fractional models fitting to the TIRFM data. **(A)** Examples of two images, each representing the same FoV, with differing temporal resolutions. **(B)** Fraction size of the fast-diffusing population (*α*), plotted against the time lag. **(C)** Mean squared displacements plotted against the time lag for a fast-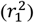 and a slow-diffusing 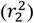 fractions for the images acquired in TIRFM with a time lag of 25 ms. **(D)** Mean squared displacements plotted against the time lag for a fast-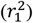 and a slow-diffusing 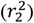 fractions for the images acquired in TIRFM with a time lag of 200 ms. Results of the fits are summarized in Table 1. To establish the values of dynamic parameters, 5 different embryos per each TIRFM imaging approach were imaged on each of the 3 different experimental days. Each datapoint is presented in the form of a mean ± s.e.m., and the 95% c.i. of the mathematical fit is shown. Shapiro-Wilk statistical test was performed to check for normality of the data set. Statistical analysis was performed using a one-way ANOVA with a Tukey’s range post-hoc test (for details of the statistical test results, see Table 2). **(E)** Cumulative probability distribution function of the GFP-C10H-Ras squared displacements calculated for the TIRFM datasets. Data points are shown in red, and their population fits in dashed green for a one-population model, in dashed blue for a two-population model, and in dashed black for a three-population model (formulae shown). Fitting of the data points to the three-population model allowed for calculation of a relative size of the subpopulations (*α*_1_ and *α*_2_) and their mean squared displacements 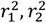 and 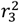. This procedure was repeated for each of the time lags used. **(F)** Mean squared displacements plotted against the time lag for a fast-diffusing, a slow-diffusing, and an immobile fraction 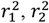, and 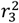, using the three-population model fit. Values of the fast-diffusing and the slow-diffusing fraction sizes, *α*_1_ and *α* _2_ are shown.

To investigate the GFP-C10H-Ras in more detail, we utilized the TIRFM data with a temporal resolution of 25 ms, presented in Fig. 5BC, and we fitted one-population (*R* ^2^ ∼ 0.912, Eq. 1), two-population (*R* ^2^ ∼ 0.982, Eq. 2), and three-population (*R* ^2^ > 0.999, Eq. 3) models to the data (Fig. 5EF, Table 1). Interestingly, the analysis for the TIRFM data exhibited that it is the three-population model that is most suitable to describe the dynamics of the H-Ras anchors imaged using this microscopy setup (Fig. 5E). Based on the three-population model for the TIRFM data, we could differentiate between a fast-diffusing, a slow-diffusing, and an immobile population. The percentage of GFP-tagged H-Ras anchors equaled 51% for the fast-diffusing fraction, and 39% for the anchors in the slow-diffusing population. Immobile molecules constituted 10% of the total GFP-C10H-Ras population (Fig. 5F). The initial diffusion coefficient *D*_0_ for the fast-diffusing population equaled 1.15 ± 0.03 μm^2^ s^-1^, and 0.306 ± 0.045 μm^2^ s^-1^ for the slow-diffusing one. The corresponding confinement area size *L* equaled 581 ± 8 nm and 286 ± 16 nm for the fast-diffusing and the slow-diffusing population, respectively. Consequently, fitting the three-population model to the TIRFM data not only showed the occurrence of three populations, but also confirmed that the fastest population of H-Ras anchors is not detected in the 2PEFM setup. The dynamic parameters of the two populations acquired by imaging GFP-C10H-Ras with the 2PEFM setup (Fig. 4CD) mostly corresponded to the slow-diffusing and immobile fraction observed in the TIRFM setup with the time lag of 25 ms (Fig. 5F).

### Analysis of the excitation laser power impact on the GFP-C10H-Ras mobility pattern

To study a possible effect of the excitation laser power on the observed mobility patterns, we used the TIRFM setup with a time lag of 25 ms and imaged GFP-C10H-Ras in zebrafish embryos with increasing excitation laser power, i.e., 10%, 30%, and 50% of the setup’s maximal power, which measured at the sample plane equaled ∼40 mW (with the illumination focal plane area of 1.65 · 10^5^ cm^2^, the laser power density equaled ∼2.42 kW cm^-2^). The results of this experiment showed no significant differences as a result of the changed laser power. The size of the fast-diffusing fraction did not differ significantly between the different laser powers applied, and equaled 61.0 ± 15.7% for 10%, 60.7 ± 9.8% for 30%, and 54.8 ± 6.3% for 50% of the total laser power (Fig. 6A). Initial diffusion coefficients *D*_0_ for the fast-diffusing fraction did not differ either, and equaled 0.91 ± 0.14 μm^2^ s^-1^ for 50%, 1.22 ± 0.22 μm^2^ s^-1^ for 30%, and 0.76 ± 0.02 μm^2^ s^-1^ for 10% of the total laser power (Fig. 6C). Similarly, the size of the confinement area *L* for illumination with 50% (444 ± 3 nm) was similar to those imaged with 30% and 10% of the total laser power (475 ± 5 nm and 536 ± 3 nm, respectively). The initial diffusion coefficients and the sizes of the confinement areas for the slow-diffusing fractions were also similar for all of the experimental groups (Fig. 6D).

To investigate whether the different populations display different fluorescence intensities and may represent clusters comprising a different number of molecules, we plotted the intensity of selected particles against their squared displacement. We selected 30 individual GFP molecules per fraction, based on a threshold value of 0.01 μm^2^, for each excitation power. Interestingly, no significant differences in the intensities were observed between the slow- and the fast-diffusing fraction, indicating that the clusters of these fractions contain a similar number of GFP-C10H-Ras molecules. As expected, the mean intensity for the molecules imaged with 50% of the TIRFM maximal laser power was the highest and equaled 47361 ± 5645 counts for the fast-diffusing fraction and 43661 ± 5137 counts for the slow-diffusing one. For the 30% of the maximal laser power, the intensity equaled 16113 ± 1993 counts for the fast-diffusing fraction and 13847 ± 1310 counts for the slow-diffusing one, whereas for the 10% of the maximal laser power the intensity equaled 2107 ± 214 counts for the fast-diffusing fraction and 1984 ± 119 counts for the slow-diffusing one (Fig. 6B).

**FIGURE 6:**
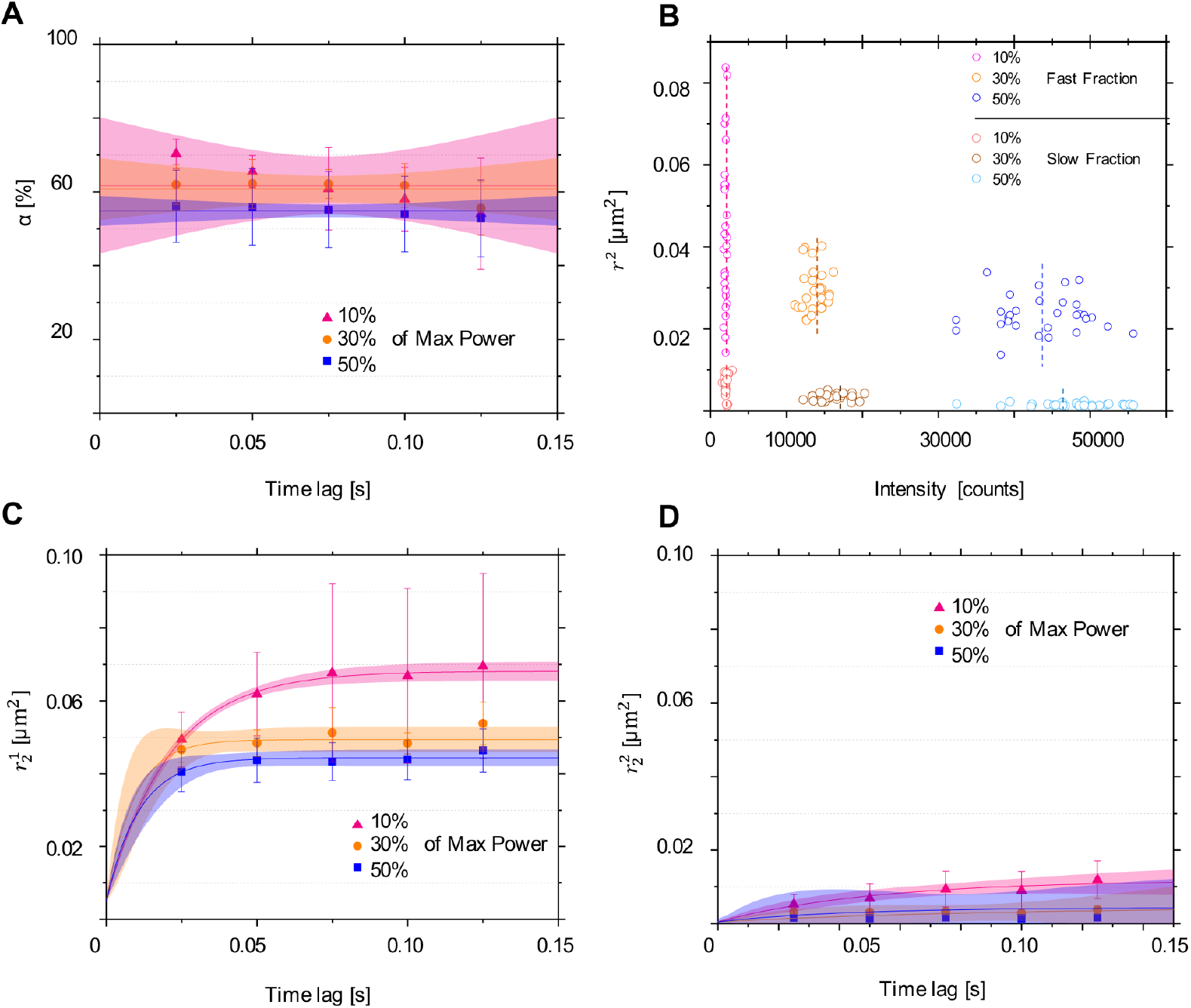
GFP-C10H-Ras mobility patterns: differences between different TIRFM excitation laser powers. **(A)** Fraction size of the fast-diffusing population (*α*), plotted against the time lag of 25 ms. **(B)** Relationship between intensity counts and squared displacement for 30 molecules selected per their fraction and applied excitation laser power. For 50% of the total laser power, the mean intensity value equaled 47361 ± 5645 for the fast-diffusing fraction and 43661 ± 5137 for the slow-diffusing one, whereas the background intensity equaled 19800 counts. For 30% of the total laser power, the mean intensity value equaled 16113 ± 1993 for the fast-diffusing fraction and 13847 ± 1310 for the slow-diffusing one, whereas the background intensity equaled 6141 ± 5645 counts. For 10% of the total laser power, the mean intensity value equaled 2107 ± 214 for the fast-diffusing fraction and 1984 ± 119 for the slow-diffusing one, whereas the background intensity equaled 534 counts. **(C)** Mean squared displacements plotted against the time lag for the fast-diffusing fraction 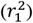. **(D)** Mean squared displacements plotted against the time lag for the slow-diffusing fraction 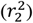. Results of the fits are summarized in Table 1. To establish the values of dynamic parameters, 5 different embryos per each TIRFM excitation laser power were imaged on each of the 3 different experimental days. Each datapoint is presented in the form of a mean ± s.e.m., and the 95% c.i. of the mathematical fit is shown. Shapiro-Wilk statistical test was performed to check for normality of the data set. Statistical analysis was performed using a one-way ANOVA with a Tukey’s range post-hoc test (for details of the statistical test results, see Table 2).

### The mobility pattern of GFP-C10H-Ras in epidermal cells of zebrafish embryos after treatment with Latrunculin B and Methyl-β-cyclodextrin

To further evaluate whether the absence of the fast subpopulation of GFP-C10H-Ras molecules is inherent to our 2PEFM-based analysis, we incubated zebrafish embryos with Latrunculin B (LatB) and Methyl-β-cyclodextrin (MBCD). In our previous study employing TIRFM, both LatB and MBCD significantly increased the size of the fast-diffusing fraction of YFP-H-Ras molecules together with its initial diffusion coefficient and confinement area size (Gora et al., 2021). Such changes, however, were not observed in the slow-diffusing fraction, making these treatments a powerful tool to distinguish between the two fractions. Thus, after treating the zebrafish embryos with LatB or MBCD, the 2PEFM was performed and the mobility patterns of GFP-C10H-Ras were analyzed. Again, we observed a slow-diffusing and an immobile fraction of molecules in all experimental groups (Fig. 7C).

**FIGURE 7:**
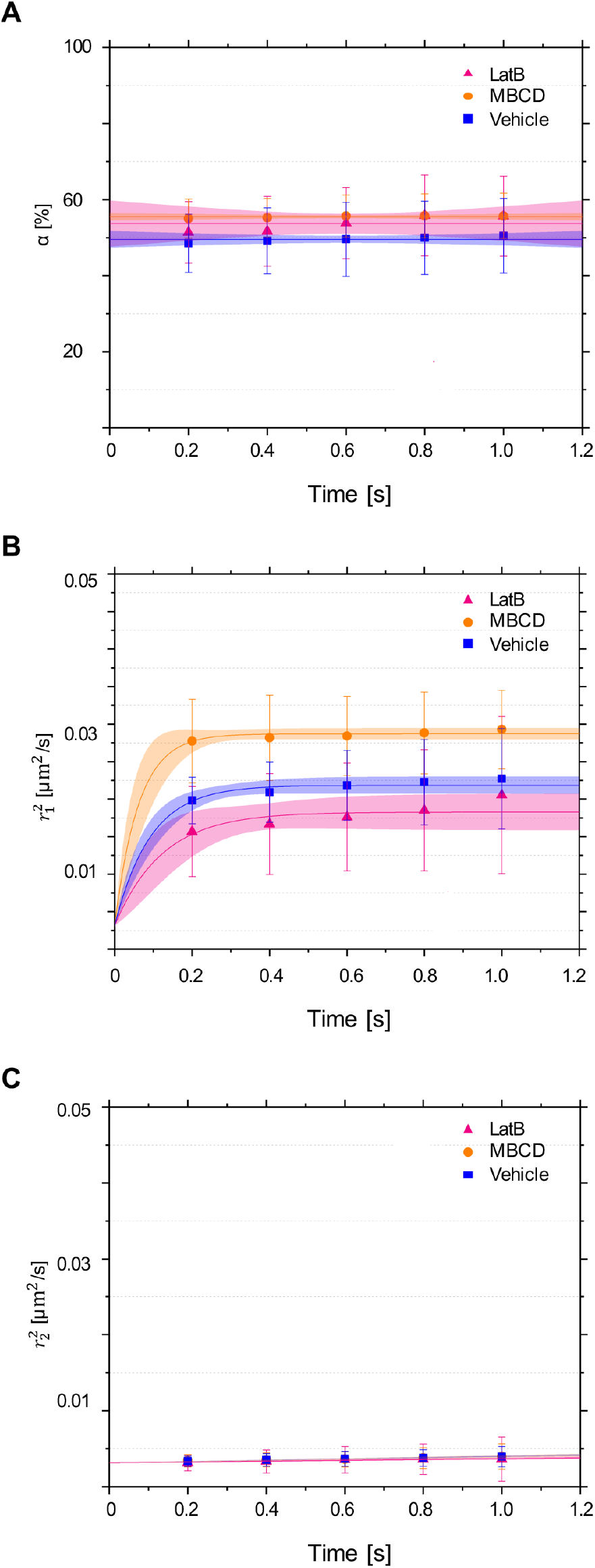
Mobility patterns of GFP-C10H-Ras in epidermal cells of the zebrafish embryos before and after treatment with Latrunculin B (LatB) and Methyl-β-cyclodextrin (MBCD). **(A, B, C)** Plots representing mobility patterns of the GFP-C10H-Ras inside the zebrafish embryos from the Tg*(bactin: GFP-C10H-Ras)*^*vu119*^ transgenic line after the treatments with a vehicle, LatB and MBCD. **(A)** Fraction size of the fast-diffusing population (*α*), plotted against the time lag. **(B)** Mean squared displacements plotted against the time lag for the fast-diffusing fraction 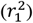. **(C)** Mean squared displacements plotted against the time lag for the slow-diffusing fraction 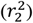. Parameters obtained upon curve fitting are presented in Table 1. To establish the values of dynamic parameters for the GFP-C10H-Ras from the transgenic line treated with vehicle, LatB and MBCD, 5 different embryos per each treatment group and imaged on each of the 3 different experimental days. Each data point is presented in the form of a mean ± s.e.m., and the 95% c.i. of the mathematical fit is shown. Shapiro-Wilk statistical test was performed to check for normality of the data set. Statistical analysis for all treatment groups was performed using a one-way ANOVA (P-value 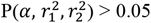 at a *t*_*lag*_ of 200 ms) with a Tukey’s range posthoc test (for details of the post-hoc test results, see Table 2).

The size of the slow-diffusing fraction did not significantly change after LatB or MBCD treatment (from 49.9 ± 0.3% in the vehicle group to 53.7 ± 0.9% and 55.5 ± 0.2% after LatB and MBCD treatment, respectively) (Fig. 7A). Additionally, treatment with LatB and MBCD did not significantly alter the initial diffusion coefficient of the slow-diffusing fraction (Fig. 7B). The size of the confinement area of the slow-diffusing fraction was not significantly changed either and equaled 237 ± 3 nm for the vehicle group, 213 ± 5 nm for the LatB treatment group, and 277 ± 2 nm for the MBCD treatment group (Fig. 7B).

These data further confirmed that, in the 2PEFM setup, we did not detect the fast-diffusing GFP-C10H-Ras fraction that was identified in our previous TIRFM experiments, and was largely affected by LatB and MBCD treatment, since the effects induced by these treatments were not observed in the current study. We, therefore, conclude that the fraction detected using the 2PEFM setup that diffused the fastest corresponds to the slow-diffusing subpopulation that we previously identified using TIRFM, which showed a similar mobility pattern and was not affected by LatB and MBCD treatment, either.

### Analysis of the single GFP-C10H-Ras trajectories based on the 2PEFM imaging

In the final part of this study, we wanted to utilize the strongly reduced photobleaching (as demonstrated in Fig. 3) to reconstruct and examine trajectories of GFP-C10H-Ras molecules from the slow-diffusing population. While the time until photobleaching of GFP or YFP molecules in SMM studies is generally in the order of milliseconds (Harms et al., 2001), the 2PEFM imaging allowed us to track the vast majority of GFP-C10H-Ras molecules for at least 15 frames with the 2PEFM integration time of 200 ms, i.e., for more than 3 seconds, which enabled us to study their dynamic behavior over a prolonged time and reconstruct their trajectories.

First, we randomly selected five cells and mapped the trajectories of the GFP-C10H-Ras particles in these cells using the R software (Fig. 8 & 9A). The trajectories were selected based on a low molecular density in their proximity, so that any possibility of crisscrossing and incorrect trace attribution was eliminated. Due to practical lack of movement, the molecules from the immobile group were not included in this analysis. In each of the selected cells, we selected five trajectories, hence 25 in total, to investigate in more detail (Fig. 8).

**FIGURE 8:**
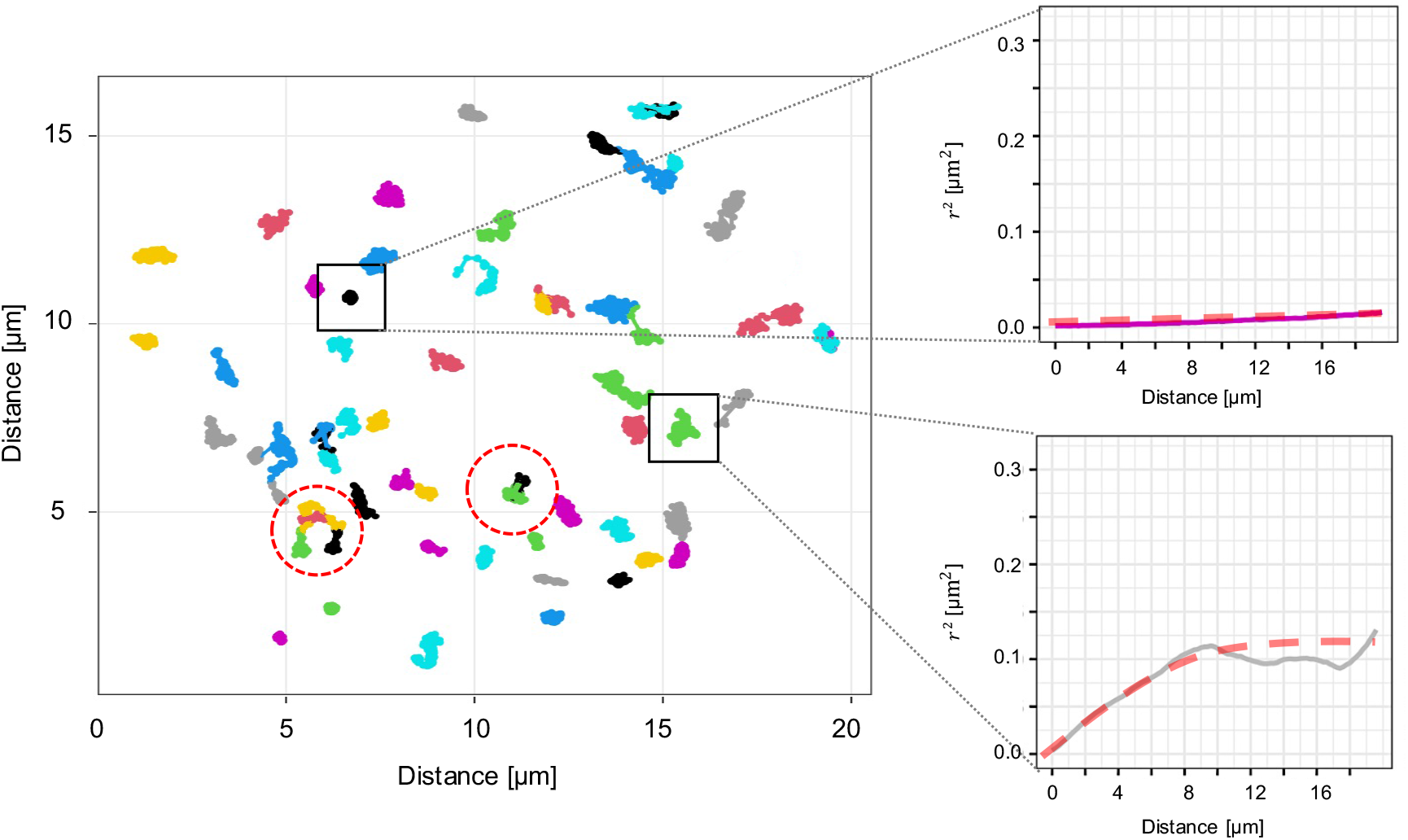
Reconstruction of H-Ras anchor molecular trajectories obtained through 2PEFM imaging. A schematic depicting the process of selection of molecules used for a single GFP-C10H-Ras trajectory analysis. Many particles are distinguished during such an analysis, but not all of those are useful for trajectory reconstruction. Particles inside black square selections denote examples of an immobile molecule and a molecule exhibiting a confined diffusion. Immobile molecules were not used in the further analysis, as they did not diffuse throughout their fluorescence lifetime. Particles inside red circular selections crisscross with each other and, therefore, also cannot be used for further analysis, since it is impossible to reliably assign a single trajectory to particles diffusing in such proximity.

The squared displacements of all 25 selected molecules were used to generate a plot presenting the mean squared displacement as a function of time, to demonstrate that this selection of molecules is representative of the entire molecular population detected, as shown in Fig. S2. The plotting of the confinement model fit the function of mean squared displacements over time revealed that the initial diffusion coefficient *D*_0_ for this population equaled 0.243 ± 0.057 μm^2^ s^-1^, while its confinement area size *L* equaled 357 ± 82 nm, values that are similar to those obtained through the one-population model fitting (Fig. S1)

Subsequently, for every one of these molecules, we plotted the squared displacement and the cumulative squared displacement over time (Fig. 9BC). What became apparent from these graphs was the fact that the diffusing GFP-C10H-Ras particles appear to switch between a diffusing state and a state that was characterized by brief bursts of increased diffusion, which may be referred to as hopping. To further analyze the alternating between these two states, we defined the diffusing state by a squared displacement between two consecutive frames smaller than 0.12 μm^2^ (i.e., *D*_0_ < 0.15 μm^2^/s), and the hopping state by a squared displacement larger than 0.12 μm^2^ (i.e., *D*_0_ > 0.15 μm^2^/s), based on the squared displacement versus time lag plots, (such as the one shown in Fig. 9, and more examples are presented in Figure S3). We validated this approach by examining mean squared displacement and confinement area size of molecular steps in the selected molecules, using the PICS software (Fig. S4). The results of this analysis exhibited that the hopping molecules diffused with a mean diffusion coefficient of 0.412 ± 0.083 μm^2^ s^-1^ in a confinement area of 1038 ± 56 nm, whereas the molecules in the diffusing mode moved with a diffusion coefficient of 0.046 ± 0.002 μm^2^ s^-1^ in confinement area of 192 ± 13 nm.

**FIGURE 9:**
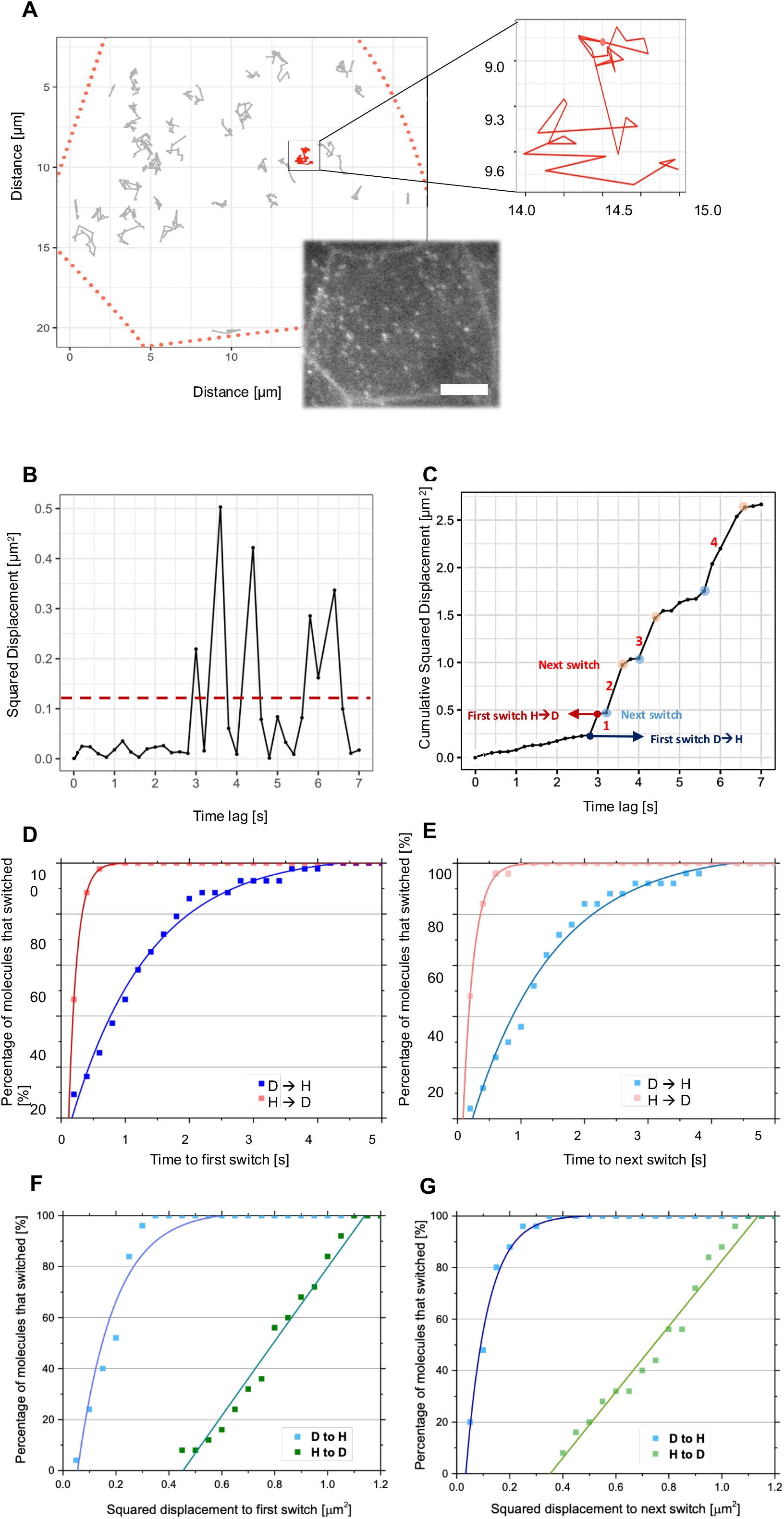
Analysis of single GFP-C10H-Ras molecular trajectories and their switches between different diffusion states. **(A)** Representative image of an epidermal cell selected for the reconstruction of the GFP-C10H-Ras molecular trajectories, and its mapped picture designed in the R software. Red dotted line signifies the cell membrane, while isolated trajectories of single molecules are shown in grey. An example of molecular trajectories is highlighted in red. *Right side:* Enlarged map of a single GFP-C10H-Ras molecule highlighted red in (A). **(B)** Representative plot of the squared displacement versus time lag for the single GFP-C10H-Ras molecule selected in (A). Periods of short-lived, increased diffusion rates are visible. The threshold for differentiation between the diffusion and hopping states is drawn in dashed red. **(C)** A plot of the cumulative squared displacements versus time lag for the single GFP-C10H-Ras molecule selected in (A). Identical short-lived periods of increased diffusion rates are visible, each of them is numbered. Molecular switches of the single GFP-C10H-Ras molecule are indicated in blue, for a switch to a hopping state, and in red, for a switch to a diffusing state. The first switch to a hopping state (D → H) is marked by blue squares, whereas the first switch to a diffusing state (H → D) is marked by red squares. The following switch to a hopping state is marked by light blue squares, whereas the first following switch to a diffusing state is marked by light red squares. **(D)** Plot of the time needed for a single GFP-C10H-Ras molecule to switch for the first time to a diffusion and a hopping state. The Y-axis represents the percentage of molecules from a population of 25 selected H-Ras anchors that committed to their first switch. **(E)** Plot of the time needed for a single GFP-C10H-Ras molecule to switch to a diffusion and a hopping state following their first switch. The Y-axis represents the percentage of molecules from a population of 25 selected H-Ras anchors that committed to their switch. **(F)** Plot of the squared displacement, in μm^2^, of a single GFP-C10H-Ras molecule when switching for the first time to a diffusion or a hopping state. The Y-axis represents the percentage of molecules from a population of 25 selected H-Ras anchors that committed to their first switch. **(G)** Plot of the squared displacement in μm^2^, of a single GFP-C10H-Ras molecule when switching to a diffusion and a hopping state following their first switch. The Y-axis represents the percentage of molecules from a population of 25 selected H-Ras anchors that committed to their switch.

The molecular trajectory analysis showed that the ratio of time spent in the diffusing versus the hopping state equaled 0.237, indicating that an average molecule spends about four times more time in the diffusing state than in the hopping state. Subsequently, we studied how rapidly a molecule switched between the diffusing and the hopping state, in order to determine the lifetimes of the two diffusion states. To this aim, we analyzed the time required for a molecule to switch from a diffusing to a hopping state and *vice versa* (Fig. 9D). It appeared that the diffusing GFP-C10H-Ras molecules remained in this state for relatively long periods, as it took approximately 1.2 s before 50% of the selected molecules had switched to a hopping state. Inversely, it took only around 0.3 s for 50% of the molecules that were hopping to switch back to the diffusing state. On average, GFP-C10H-Ras molecules resided for 2.74 ± 0.32 s in the diffusing state before switching, and only 0.24 ± 0.08 s in the hopping state. We analyzed the time required for the selected group of molecules to switch again to a diffusing or a hopping state after having already switched their diffusion state at least once (Fig. 9E), and no significant differences with the analysis of the first switches were observed, as 50% of the selected diffusing molecules switched again to a hopping state after approximately 1.2 s, and it took around 0.25 s for 50% of the hopping molecules to switch back. Ultimately, we determined the squared displacement of the selected molecules before switching from a diffusing to a hopping state and *vice versa* (Fig. 9FG). At a squared displacement of less than 0.2 μm^2^, 50% of molecules in the diffusing state switched to a hopping state (Fig. 9F). The molecules in the hopping states showed a larger displacement before returning to the diffusing state (at 0.8 μm^2^, 50% switched) and they displayed a bigger heterogeneity in the displacement before switching (Fig. 9G). Again, no differences were observed before the first and the following switches.

Taken together, in this study we focused on the slow-diffusing fraction of GFP-C10H-Ras molecules in the apical membranes of the epidermal cells in living zebrafish embryos, and show that this population of H-Ras membrane anchors occurs in two different dynamic states. The majority of the time it is found in a diffusing state, which is interrupted by brief periods of hopping. This finding indicates that the mobility pattern of this population of GFP-C10H-Ras molecules can be described as ‘hop’ diffusion through the plasma membrane.

## DISCUSSION

In the present study, we have applied multifocal 2PEFM to detect individual GFP-C10H-Ras molecules in epidermal cells of living zebrafish embryos. In previous studies using TIRF microscopy, we detected a fast and a slow-diffusing fraction of H-Ras molecules, whereas in this study we exclusively observed the slow-diffusing population of molecules (Gora et al., 2021; Schaaf et al., 2009). This population of GFP-C10H-Ras molecules was found to occur in clusters within the plasma membrane, and to alternate between two diffusion modes, leading to a mobility pattern that can be described as hop diffusion.

In previous SMM studies on the mobility of H-Ras in zebrafish embryos and cultured cells, it was shown that there are two main subpopulations of fluorescently labelled H-Ras or C10H-Ras molecules in the plasma membrane: one that is diffusing at a fast rate and one that is diffusing at a slow rate (Gora et al., 2021; Lommerse et al., 2004; Schaaf et al., 2009). The fast subpopulation of H-Ras molecules shows confinement to a domain of 400-500 nm and is suggested to form nanoclusters that are located in small (< 15 nm) lipid rafts (Plowman et al., 2005; Prior et al., 2001; Prior et al., 2003). These rafts are considered to be dependent on the presence of cholesterol, since their mobility is altered by LatB or MBCD treatment (Garcia-Parajo et al., 2014; Lingwood and Simons, 2010; Nickels et al., 2015; Zhou and Hancock, 2015). The wild type H-Ras protein shows a similar fast-diffusing subpopulation. However, a constitutively active mutant of H-Ras displays a fast-diffusing subpopulation that is confined to a larger domain (∼600 nm) and its mobility is not affected by LatB or MBCD treatment. These data indicate that upon activation of H-Ras its affinity for specific plasma membrane microdomains with different lipid compositions is altered (Hancock and Parton, 2005; Plowman et al., 2005; Prior et al., 2001; Prior et al., 2003; Zhou et al., 2018). In addition, activated GTP-loaded H-Ras molecules have an increased probability to form clusters that are transiently (< 1 s) immobilized (Murakoshi et al., 2004). These clusters are cholesterol-independent, i.e., their mobility is not affected by LatB or MBCD treatment, and are considered to be the sites where I the H-Ras interacts with cytoplasmic proteins such as Galectin-1 and Sur-8, and where active signaling occurs (Belanis et al., 2008; Hancock and Parton, 2005; Herrero et al., 2016; Li et al., 2000; Shalom-Feuerstein et al., 2008; Zhou et al., 2018).

To determine which previously observed H-Ras populations were represented by the molecules observed using the multifocal 2PEFM setup, we compared their dynamic behavior with that of GFP-C10H-Ras molecules found in previous studies. We observed that the mobility of the molecules detected in the 2PEFM data corresponded most closely to the slow-diffusing fraction observed in the previous studies (Gora et al., 2021; Lommerse et al., 2004; Schaaf et al., 2009). We detected the slow-diffusing population of GFP-C10H-Ras proteins, which was previously shown to be not affected by LatB and MBCD treatment (Gora et al., 2021). Indeed, these treatments did not significantly change the mobility pattern of the detected particles. Thus, based on its dynamic behavior and the insensitivity of this behavior to LatB and MBCD treatment, we conclude that the population of GFP-C10H-Ras molecules observed in our 2PEFM study represents the slow-diffusing fraction of molecules that were observed in previous studies. The main reason for the lack of detection of the fast subpopulation of GFP-C10H-Ras molecules was shown to be the relatively large integration time used in the 2PEFM experiments (200 ms). During these relatively long illumination and detection periods, the signal of the fast-diffusing molecules is spread over the relatively large area in which they move during this integration time, causing a phenomenon called motion blur (Phillips et al., 2020; Travers et al., 2020). Our study, therefore, focuses on the mobility pattern of the slow-diffusing population of GFP-C10H-Ras molecules, which we have been able to investigate in great detail.

Based on the analyzed mobility patterns of GFP-C10H-Ras over short periods (< 1s), the molecules of the slow-diffusing population detected in the present study could, in turn, be subdivided into two fractions: a slow-diffusing population that shows confinement of 253 ± 3 nm, and one that is virtually immobile, with a confinement zone of 94 ± 5 nm. Including the fast-diffusing population of molecules that is undetectable in 2PEFM, this implies that there are three fractions of GFP-C10H-Ras molecules based on their dynamic behavior: a fast- and a slow-diffusing and an immobile population. This was confirmed when we fitted our TIRFM data (obtained with a time lag of 25 ms) to a three-population model. However, we also showed that this bulk analysis over such a short time by plotting mean squared displacements over time does not reveal the complexity of the single-molecule mobility patterns. Interestingly, our multifocal 2PEFM approach allowed us to follow the single particles for relatively long periods, which enabled us to analyze molecular trajectories of single GFP-C10H-Ras particles in the slow-diffusing population. Using this analysis, we identified two different dynamic states in which the slow-diffusing molecules occur: a state of diffusion and a state of hopping. The immobile fraction has not been included in the analysis, as the particles in this subpopulation hardly ever diffused, which is reflected by their initial diffusion coefficient of (0.080 ± 0.010) × 10^−2^ μm^2^ s^-1^.

By examining the switching of the H-Ras anchor between the two different states, we demonstrated that these anchors spend a relatively long time in the diffusion state, since the total time spent in this state was approximately four times longer than the time spent in the hopping state. Based on these findings we concluded that the observed slow-diffusing population of H-Ras anchors exhibit anomalous diffusion patterns, characterized by short-lived bursts of fast-speed diffusion, generally referred to as molecular hopping. It has been reported that many signaling proteins experience two-dimensional hop diffusion on the membrane due to the complex and packed structure of the plasma membrane (Campagnola et al., 2015; Umemura et al., 2008; Yasui et al., 2014). The plasma membrane is divided into microdomains by the actin-based membrane skeleton, which is closely associated with the cytoplasmic surface of the plasma membrane (Morone et al., 2008). Many transmembrane proteins collide with this membrane skeleton, which induces temporary confinement of the transmembrane proteins in the membrane-skeleton meshwork. Moreover, in the vicinity of immobilized molecules located in the membrane, the movement of any other particles is extremely limited, also because of the hydrodynamic dragging effects of the transmembrane proteins, which further suppress the available membrane space (Murase et al., 2004). In addition, it has been proved that interactions with many cytoplasmic proteins, such as Galectin-1 and Sur-8, are involved in the immobilization of the H-Ras clusters, which are considered to be the sites where active signaling occurs (Belanis et al., 2008; Hancock and Parton, 2005; Herrero et al., 2016; Li et al., 2000; Shalom-Feuerstein et al., 2008; Zhou et al., 2018). With all these obstacles, membrane proteins and lipids often hop from one microdomain to an adjacent one, especially when thermal fluctuations of the membrane and the actin meshwork associated with it create sufficient space between them to enable the passage of integral membrane proteins, or when an actin filament is temporarily severed (Suzuki et al., 2005).

Such a temporary nature of protein-membrane interactions enables a tight temporal regulation of signal transduction processes. It has also been proposed that molecular hopping might be critical in the search for target molecules in eukaryotic cells. A straightforward consequence of membrane hopping is that a molecule remains in its immediate vicinity for a short time and then jumps to a location that is further away than expected from two-dimensional diffusion. In such a way, the search process is allowed to explore larger areas and what allows proteins to bypass diffusion barriers that may be present in the membrane (Lemmon, 2008). As this process seems to be governed predominantly by membrane-protein associations, including electrostatic interactions, it is highly possible that the H-Ras anchor, being the domain that is responsible for attachment of the H-Ras protein to the cell phospholipid membrane, undergoes this type of anomalous diffusion.

The multifocal 2PEFM technology we used in this study induced strongly reduced photobleaching of GFP molecules compared to conventional, one photon excitation-based SMM technologies, which enabled us to visualize the diffusing proteins for prolonged periods. There may be three reasons for this reduced photobleaching. First, the multifocal 2PEFM setup used, combined with a low-noise sCMOS camera allowed for a low excitation power, being typically of 1.4 mW per focus while maintaining a relatively high signal-to-noise ratio (SNR > 10). This gentle illumination must have contributed to the low bleaching rate of GFP, since this bleaching rate in 2PEFM has been shown to be highly dependent on the laser power, even more than in one-photon excitation. The log-log plot of the excitation power versus the photobleaching rate for one-photon excitation of fluorescein increased with a slope of ∼1, whereas this slope was ∼3 for two-photon excitation (Graham et al., 2015; Patterson and Piston, 2000). Second, since previous research has indicated that the faster photobleaching processes occur at a fluorophore’s higher energetic states, the high fluorophore stability of the GFP molecules that we report may suggest that photobleaching is reduced because fluorophores are excited to their lowest excited energy state (Dittrich and Schwille, 2001; Patterson and Piston, 2000). Third, the high photostability of GFP in our 2PEFM setup might have also been the result of the pulsed excitation mode, due to the use of a pulsed laser combined with multifocal excitation. Many fluorescent molecules end up in a relatively long-lived dark state after excitation, and in this dark state the fluorescent molecules experience a highly increased chance to undergo an irreversible photobleaching reaction. By a rapid and pulsed illumination, the fluorophores residing in their dark (triplet) state have sufficient time between subsequent illuminations to undergo relaxation back to their ground (singlet) state, leading to the overall increase in photostability and fluorescent signal yield (Donnert et al., 2007; Donnert et al., 2009). On top of the fluorophore stability, non-fluorescent absorption is known to cause damage to biological tissues, and the 2PEFM technology with low phototoxicity is, therefore, highly suitable for studies in *in vivo* systems such as zebrafish embryos (Eggeling et al., 1998; Niesner et al., 2007).

Finally, we studied the nature of the slow-diffusing population of GFP-C10H-Ras particles by determining the intensity of the fluorescence intensity peaks and their photobleaching profile over time. It became apparent that the molecules occur in clusters, since we detected individual fluorescent molecules as well as signals that could be attributed to multiple GFP-C10HRas proteins. It has previously been shown that C10H-Ras, as well as the full-width H-Ras, often reside in small clusters with a radius varying from 40 to 180 nm (Zhou and Hancock, 2021). Moreover, it has been reported that C10H-Ras localizes to and clusters in cholesterol-rich lipid rafts, however, upon their disruption, disperses over the surface of the plasma membrane, rather than driving associations with other membrane microdomains. These findings are consistent with our data and support a biodynamic model where full-length H-Ras has an affinity for at least two plasma membrane microdomains, lipid rafts and a cholesterol-independent non-raft microdomain, and is in constant equilibrium between them (Murakoshi et al., 2004; Prior et al., 2003).

In conclusion, in the present study, we showed that 2PEFM is a powerful microscopy technique for imaging single molecules *in vivo*. With this technique, we could detect the slow-diffusing population of GFP-C10H-Ras in the membranes of epidermal cells in living zebrafish embryos. We revealed that this population occurs in clusters and displays hop diffusion through the membrane, in which periods of a slow diffusion are interspersed with brief periods of molecular hopping.

## MATERIALS AND METHODS

### Zebrafish

Zebrafish larvae (*Danio rerio*) from the transgenic line *Tg(bactin: GFP-C10H-Ras)*^*vu119*^ were maintained according to standard protocols (http://ZFIN.org), and exposed to a 14h light and 10h dark diurnal cycle at 28°C. Fertilization was performed by natural spawning at the beginning of the light period. Eggs were collected and raised in egg water (60 μg/ml Instant Ocean Sea salts, Cincinnati, OH, USA) at 28°C. The eggs developed in an incubator at 28°C until 2 days post-fertilization (2 dpf). The viability and development of the eggs were checked daily using fluorescence stereo- or confocal microscopy. All experiments performed on living zebrafish embryos were done in compliance with the directives of the local animal welfare committee of Leiden University.

### Fluorescence stereomicroscopy

To screen zebrafish embryos for optimal expression levels of the GFP-C10H-Ras, a Leica M205FA fluorescence stereomicroscope (Leica Microsystems) was used. Images of the zebrafish embryos were taken using a Leica DFC 345FX camera.

### Treatment of zebrafish embryos with Latrunculin B (LatB) and Methyl-β-cyclodextrin (MBCD)

Inhibition of actin polymerization with LatB was induced using a protocol described previously (Kugler et al., 2019). LatB (Sigma-Aldrich, St. Louis, MO, USA) was dissolved in 96% ethanol to a 500 μM stock concentration. At 48 hpf, dechorionated embryos were treated with 100 nM LatB in egg water for 1 hour. A control group was treated with a diluted vehicle (0.02% ethanol in egg water). Treatment with MBCD was based on protocols described before (Bello-Perez et al., 2020; Silva et al., 2019). MBCD (Sigma-Aldrich, St. Louis, MO, USA) was dissolved in PBS (pH 7.4) to a stock concentration of 400 nM. At 48 hpf, dechorionated embryos were treated with MBCD at a final concentration of 40 nM in egg water for 1 hour, A control group was treated with diluted vehicle (10x diluted PBS in egg water). After the LatB or MBCD treatment, zebrafish were immediately transferred and imaged under the 2PEFM setup.

### Sample preparation and mounting

Glass coverslips were washed with 99% ethanol (twice), HPLC-grade water (twice), KOH (1M, twice), and acetone (99%, thrice). Each wash was followed by a 30-minute-long sonication period at 50°C. The coverslips were then stored in 99% acetone. Prior to the mounting of the zebrafish embryos, the glass coverslips were coated with 50 μg ml^-1^ of poly-L-lysine (Sigma-Aldrich, St. Louis, MO, USA) for 5 minutes, followed by a double wash with deionized water and drying with nitrogen gas. Two-day-old zebrafish embryos were equilibrated at room temperature for an hour, anaesthetized with 0.02% aminobenzoic acid ethyl ester (tricaine, Sigma-Aldrich St. Louis, MO, USA), and dechorionated using tweezers. Subsequently, a single zebrafish embryo was placed on a coverslip with a lateral side against the surface, while excess water was aspirated. The tail of the embryo was pressed against the coverslip by a thin agarose sheet (2%, thickness 0.75 mm). A drop of egg water was added to cover the rest of the embryo’s body. The coverslip with the embryo was placed into the microscope holder, which was then inserted into the imaging chamber.

### Two-Photon Excitation Fluorescence Microscopy (2PEFM)

For 2PEFM, a tunable near-IR Ti: Sa laser (Chameleon Ultra; Coherent, Santa Clara, CA, USA) was coupled into a home-built two-photon multifocal microscope. A diffractive optical element (HOLOEYE Photonics AG, Berlin, Germany) diffracted the laser beam into an array of 10 × 10 beamlets. A fast-scanning mirror (FSM-300-1; Newport, Irvine, CA, USA) scanned the beamlets across the excitation plane. The laser beams were focused using a 60x oil immersion objective (CF160 Apochromat TIRF 60XC, NA 1.49; Nikon, Tokyo, Japan) mounted on a piezo-stage (P-726 PIFOC; Physik Instrumente, Karlsruhe, Germany), illuminating an area of 40 μm x 40 μm. Two-photon luminescence was collected by the same objective, filtered with a dichroic mirror (700dcxr; Semrock, Rochester, NY, USA) and two short-pass filters (FF01-720-SP and FF01-750-SP; Semrock), and focused on a 2048 × 2048 pixel back-illuminated sCMOS camera (Prime BSI; Teledyne Photometrics, Tucson, AZ, USA). Additional band-pass filters, mounted on a motorized fast-change filter wheel (FW103H/M; Thorlabs, Newton, NJ, USA), were positioned in front of the camera. Using custom-made LabVIEW software (National Instruments, Austin, TX, USA), the scanning mirror, focusing stepper motors and camera were controlled synchronously. For spiral scanning, the fast-scanning mirror (FSM-300-1, Newport) was driven by an Archimedean spiral to rapidly scan the beams producing a homogeneously illuminated wide-field. A single period of the spiral scan took 200 ms and was synchronized with the camera integration time, resulting in a temporal resolution of 200 ms. The excitation laser power at the sample surface equaled 0.1497 mW at the beginning of a time-lapse, and decreased to 0.1425 mW at the end of the recording. Every embryo was imaged for at least 2 minutes, which equaled a minimum of 600 frames in a single time-lapse. All 2PEFM measurements were performed at room temperature.

### Total internal reflection fluorescence microscopy (TIRFM)

For TIRFM, a custom-made microscope was used with a 100x oil-immersion objective (NA 1.45, Nikon, Tokyo, Japan). Excitation was performed using a 515 nm laser (iChrome MLE, Toptica Photonics, Germany), the field of view was set to a 100 × 100 pixels region with a pixel size of 166 nm, and the laser power equaled 20% of the maximal laser power (40 mW). The incident laser beam was set at the critical angle against the coverslip-water interface, thus being totally reflected and creating the evanescent wave for the excitation of fluorophores close to the coverslip-sample interface. Emission light was filtered using a long-pass filter (ET5701p, Chroma Technology, VT, USA), and image sequences were collected using an on-chip multiplication gain CCD camera (model 512B, Cascade, Roper Scientific, Tucson, AZ, USA). Each image sequence contained 1200 frames separated by a 25 or 200 ms time lag, resulting in a total acquisition time of 30 and 240 seconds, respectively. All TIRFM measurements were performed at room temperature.

### Analysis of protein diffusion patterns

Fluorescence intensity signals corresponding to GFP molecules were fitted to a two-dimensional Gaussian surface, using custom-developed software (Groeneweg et al., 2014; Harms et al., 2001; Lommerse et al., 2004; Lommerse et al., 2005; Schütz et al., 1997a). The software, in the form of a code written in the MATLAB programming environment (The MathWorks, Natick, MA, USA), can be obtained by directly contacting Thomas Schmidt (Leiden Institute of Physics, Leiden University). The location of a particle was defined as the position of the maximum of the Gaussian curve. The Gaussian curve fitting provided information on the intensity and full width at half maximum (FWHM) for the localized GFP peaks, which allowed for plotting the intensity distributions of the GFP peaks for each frame in a time-lapse. The positional accuracy *dx* of the peak localization equaled approximately 28 nm (Groeneweg et al., 2014; Schütz et al., 1997).

To identify subpopulations of molecules, their diffusion coefficients, and the sizes of their confinement area, the Particle Image Correlation Spectroscopy (PICS) software was used to analyze distributions of squared displacements for individual time lags. The PICS software, a code written in the MATLAB programming environment (The MathWorks, Natick, MA, USA), has previously been described and validated (Semrau and Schmidt, 2007), and can be obtained can by directly contacting Thomas Schmidt or Stefan Semrau (Leiden Institute of Physics, Leiden University). In PICS analysis, individual particles are not tracked, but correlations between the location of molecules in consecutive frames are determined. A multistep analysis was performed for each image sequence acquired, yielding information for five different time lags of 200, 400, 600, 800, and 1000 ms. This way, cumulative probability distributions of the squared displacements were generated for each time lag and fitted to one-, two- or three-population models. The one-population model is described by the equation:

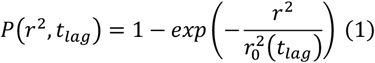

which describes the probability that a particle exhibiting Brownian motion at the arbitrary origin is found within a circle of a radius *r* at the time lag *t* _*lag*_ and its mean square displacement equals 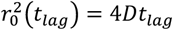. However, when a two-population model is used, Equation 1 is transformed into Equation 2:

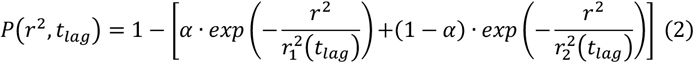

and, when a three-population model is used, into Equation 3:

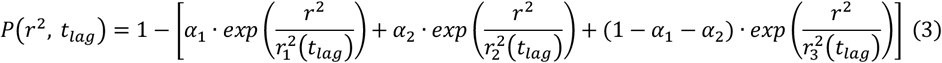

where the mean squared displacements of the populations are denoted by 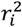, and their relative sizes by *α*_(*i*)_ (Lommerse et al., 004; Schaaf et al., 2009). In the two-population model, the fraction size *α* represents a percentage of the fast-diffusing H-Ras anchor molecules in the total population. The mean squared displacements of the fast- and slow-diffusing fractions in this model are denoted by 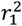 and 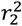, respectively. The three-population model accounts for an additional fraction, denoted as 1 − (*α*_1_+ *α*_2_), and is characterized by the mean squared displacement, 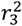.

To examine whether any of these populations confine to specific areas, the values of 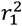 and 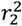, were plotted against the time lag (using OriginLab software, OriginLab Corporation, MA, USA). The positional accuracy *dx* led to a constant offset in *r*^*2*^ of 4 · (*dx*)^2^, which, in our case, equaled 0.0031 μm^2^. The plots were fitted either to a free Brownian diffusion model, with a diffusion coefficient *D* determined by the equation

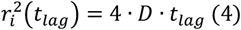

or to a confined diffusion model described by the equation:

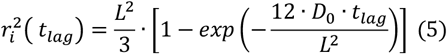

in which the molecules move freely with an initial diffusion coefficient *D*_0_, but are confined to an area with impermeable barriers, described by a square of a side length *L*.

### Analysis of photobleaching

Using an ImageJ plug-in, TrackMate (Tinevez et al., 2017), every fluorescence intensity peak, identified by the Gaussian fitting described above, was followed over time. Throughout this tracing, the threshold on the particle diffusion rate was set to 1.10 ± 0.15 μm^2^/s, based on previous data on the H-Ras anchor mobility rates obtained in living zebrafish embryos (Gora et al., 2021). In every frame, the particle was attributed to a circular area with a radius of 5 pixels. For each particle, the intensity was measured, frame-by-frame, by determining the maximal pixel intensity found within the circular selection. The intensities of molecules with differing initial intensities were plotted against time to study the step-wise decrease in fluorescence intensity as a result of photobleaching. The mean intensity values of all particles identified within a frame and the background intensity were determined as well (based on a sample of five zebrafish cells, each one imaged on a different experimental day). Upon loss of the signal (i.e., when the intensity was similar to the value of the background signal), the area of 3 × 3 μm that surrounded the last position of a particle was followed for the next 20 frames to determine if this loss was irreversible.

### Analysis of GFP-C10H-Ras trajectories

The trajectories of GFP-C10H-Ras particles in the zebrafish epidermal cells were analyzed using the ImageJ plug-in TrackMate and the R Project software (The R Foundation, Vienna, Austria). To eliminate the impact of crisscrossing among particles, traces present in high-density areas were excluded. In addition, immobile particles were excluded. For the included traces, squared displacements and cumulative squared displacements were plotted versus the time lag and the distance, represented by squared displacement, for each step traversed by a particle until its photobleaching (OriginLab, OriginLab Corporation, MA, USA). Switches between two distinct diffusion states exhibited by the particle were then identified based on a threshold value for the squared displacement between the two consecutive frames. Following that, the lifetime of a particle in either of the two diffusing states was analyzed by generating a distribution plot of the time lengths until the first switch. Additionally, to examine whether the first diffusion switch of a molecule induced a more sustained mobility change, a similar plot was generated for all subsequent switching events as well.

### Experimental design

Five independent experiments, performed on five different days, were conducted for the transgenic zebrafish line expressing the GFP-C10H-Ras construct. In every experiment, five different zebrafish embryos were selected for the 2PEFM imaging. In each of the selected embryos, at least six separate areas within the zebrafish tailfin were imaged one time, with each time lapse comprising at least 50 consecutive frames (Fig. 2). To study the influence of zebrafish treatment with LatB and MBCD on the GFP-C10H-Ras dynamics, a similar design was implemented, with five individual embryos used per vehicle, LatB, and MBCD treatment groups, each imaged on three different experimental days (Fig. 7). Trajectories of isolated particles, localized in the cell areas not affected by crisscrossing were reconstructed in five randomly selected zebrafish cells, each one imaged on a different experimental day. Ultimately, 25 trajectories, five per every zebrafish cell, were selected to examine the switching between different diffusion states (Fig. 9).

To compare 2PEFM with the TIRFM techniques, three independent TIRFM experiments on three individual days were performed, in which five zebrafish embryos per group were used, and six independent areas per embryo were imaged, with a time lag of 25 and 200 ms. Every area was imaged once with each time lapse comprising at least 50 consecutive frames (Fig. 5). Time until photobleaching was examined using a module in the PICS software, using data from three zebrafish cell areas imaged on three individual days. The number of particles and trajectories included in this analysis are shown in the Figure 3. Subsequently, to investigate the impact of the excitation laser power on the single-particle dynamics in the TIRFM setup, three independent experiments on three individual days were done, with six independent areas in each of the five zebrafish embryos imaged. The experimental groups were exposed to 50%, 30%, and 10% of the maximal excitation laser power, and a time lag of 200 ms. Following that, 30 particles per fast- and slow-diffusing fractions in all of the experimental groups were selected to test a potential correlation between their intensity count and squared displacement values (Fig. 6). The selection of the particles was performed in TrackMate based on their squared displacement values. The threshold of the mean squared displacement values used for differentiating between the two fractions was set to 0.01 μm^2^, hence every molecule above this value was assigned to the fast-diffusing subpopulation, whereas every molecule below to the slow-diffusing subpopulation.

### Statistical Analysis

Values of the different population sizes and their squared displacements were averaged per experimental day, and their standard errors of the mean (s.e.m.) were calculated. Statistical analysis of these data was performed for the experimental time lag of 200 ms by comparing: (a) results obtained for the transgenic embryos expressing GFP-C10H-Ras, with and without LatB or MBCD treatment; (b) results obtained for transgenic embryos expressing GFP-C10H-Ras imaged using the 2PEFM with the time lag of 200 ms and the TIRFM with the time lags of 25 and 200 ms; (c) results obtained for the transgenic embryos expressing GFP-C10H-Ras, imaged under the TIRFM with 50%, 30%, and 10% of the total excitation laser power. Initial diffusion coefficients and confinement area sizes were obtained by averaging values of mean square displacements per time lag, plotting them against time and fitting a confined diffusion model yielding these parameters. This was done for each treatment, microscopy technique, and the percentage of the TIRFM total excitation laser power. In each case, a Shapiro-Wilk statistical test was performed to check if data were normally distributed. The significance of the results was analyzed using a Student’s t-test for a comparison of means between two, normally distributed, groups. In case multiple groups were compared, a one-way ANOVA was used with a Tukey range test for post hoc analysis.

## COMPETING INTERESTS

The authors declare no competing financial interests.

## FUNDING

R.J.G. was funded by Marie Sklodowska-Curie ITN project ‘ImageInLife’ (Grant Agreement No. 721537)

## AUTHOR CONTRIBUTIONS

Conceptualization: M.J.M.S.; Methodology: R.J.G., R.V, M.J.M.S.; Software: R.J.G., R.V., J.N.; Validation: R.J.G., R.V., S.J., M.J.M.S.; Formal analysis: R.J.G., R.V., S.J.; Investigation: R.J.G., R.V., S.J; Resources: J.N., M.J.M.S.; Data curation: R.J.G., S.J., .N., M.J.M.S.; Writing - original draft: R.J.G., M.J.M.S.; Writing - review & editing: R.J.G., J.N., M.J.M.S.; Supervision: J.N., M.J.M.S.; Project administration: M.J.M.S.; Funding acquisition: M.J.M.S.

## SUPPLEMENTARY FIGURES

**FIGURE S1:**
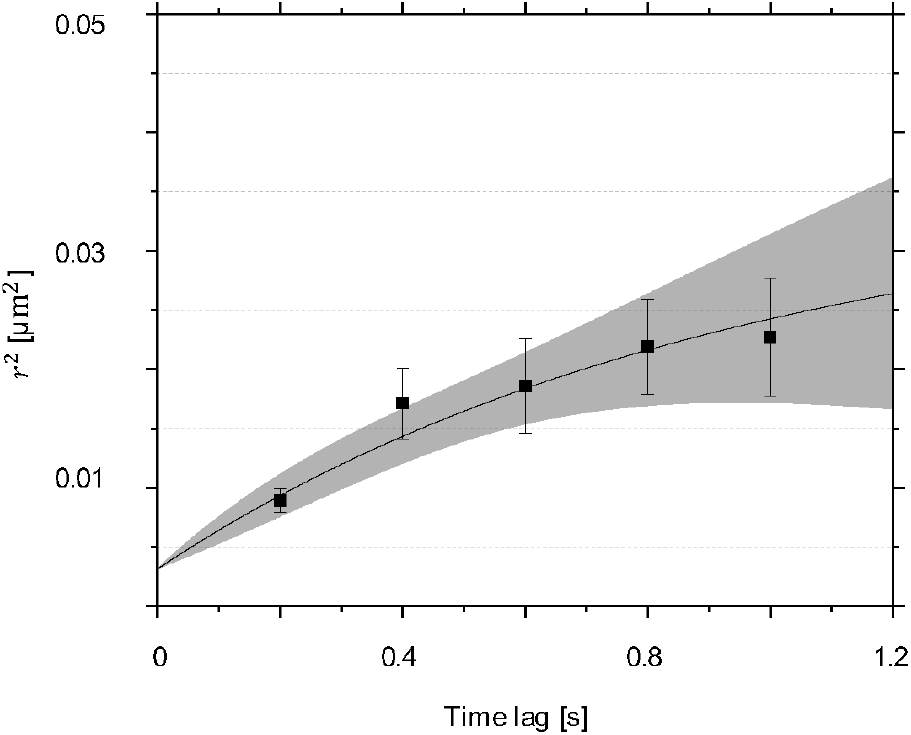
GFP-C10H-Ras mobility patterns: results of one-fractional model fitting to 2PEFM data. Fitting of the data points to the one-population model allowed for calculation of the mean squared displacements of this population (*r*^2^). This procedure was repeated for each of the time lags used. Mean squared displacements plotted against the time lag, using the one-population model fit. The initial diffusion coefficient *D*_0_ of this single population equaled 0.116 ± 0.011 μm^2^ s^-1^ and its confinement area size *L* equaled 302 ± 54 nm Each data point is presented in the form of a mean ± s.e.m., and the 95% c.i. of the mathematical fit is shown.

**FIGURE S2:**
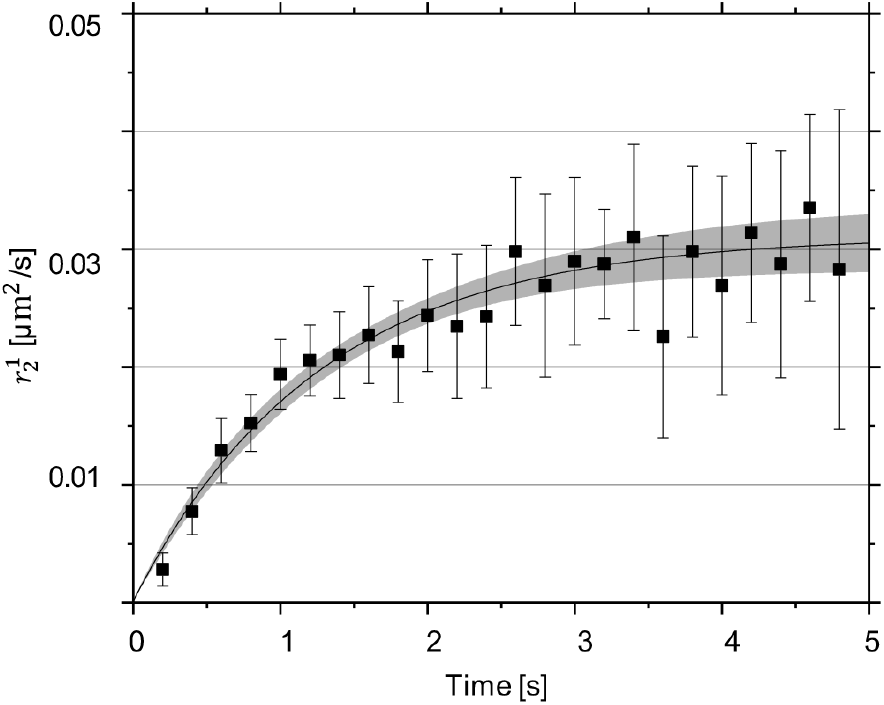
GFP-C10H-Ras mobility patterns: mean squared displacement of the molecular trajectories reconstructed for 25 selected particles. The squared displacements of all 25 selected molecules were used to generate a plot presenting the mean squared displacement as a function of time, to demonstrate that this selection of molecules is representative of the entire molecular. The plotting of the confinement model fit the function of mean squared displacements over time revealed that the initial diffusion coefficient *D*_0_ for this population equaled 0.243 ± 0.057 μm^2^ s^-1^, while its confinement area size *L* equaled 357 ± 82 nm

**FIGURE S3:**
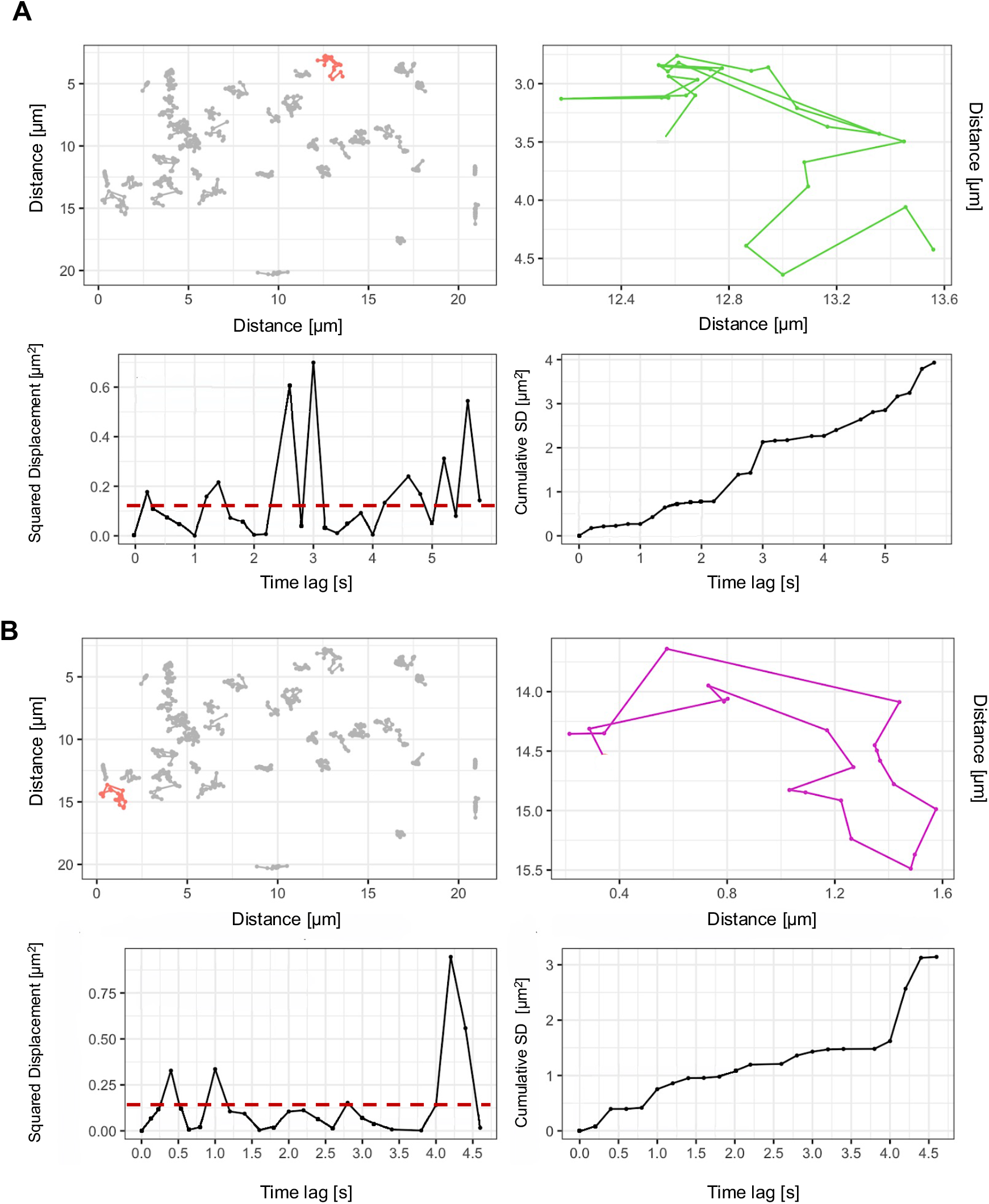
Representative images of GFP-C10H-Ras molecular trajectories. **(AB)** Representative images of two molecular trajectories used for the hop diffusion analysis. *Upper left in* **(AB)**: Inside of the epidermal cell selected for the reconstruction of the GFP-C10H-Ras molecular trajectories. Isolated trajectories of GFP molecules are shown in grey. An enlarged example of molecular trajectories is highlighted in red. *Upper right in* **(AB)**: Enlarged map of a single GFP-C10H-Ras molecule highlighted red in the upper left panel. *Lower left in* **(AB)**: A plot of the squared displacement versus time lag for the GFP-C10H-Ras molecule selected in the upper left panel. Periods of short-lived, increased diffusion rates are visible. The threshold for differentiation between the diffusion and hopping states is drawn in dashed red. *Lower right in* **(AB)**: A plot of the cumulative squared displacements versus time lag for the single GFP-C10H-Ras molecule selected in the upper left panel. Identical short-lived periods of increased diffusion rates are visible.

**FIGURE S4:**
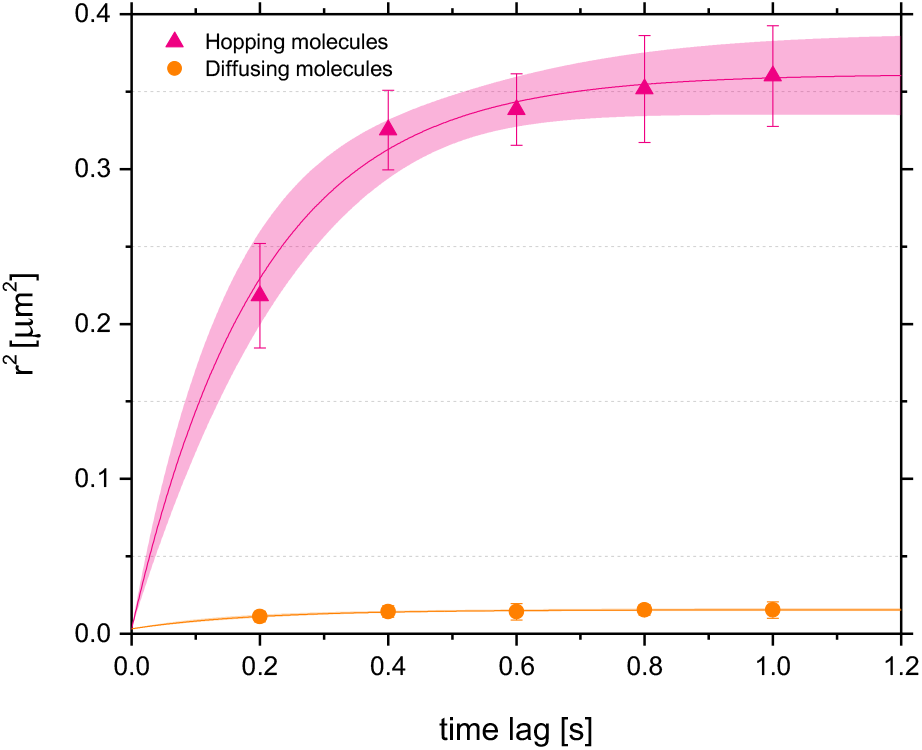
Mobility patterns of GFP-C10H-Ras molecules selected for the trajectory analysis exhibit hop diffusion. Mean squared displacements plotted against the time lag for the hopping and diffusing molecules is shown. To establish the values of dynamic parameters in these molecular fractions, trajectories of 25 fluorescent molecules, 5 per independent cell area, were selected. Each data point is presented in the form of a mean ± s.e.m., and the 95% c.i. of the mathematical fit is shown. Shapiro-Wilk statistical test was performed to check for the normality of the data set.

**TABLE S1:** Summary of the GFP-C10H-Ras mobility patterns acquired using the PICS for the molecules selected for the trajectory analysis.

| Mobility patterns of molecules selected for the trajectory analysis | $D_0$ [$\mu\text{m}^2 \text{s}^{-1}$] | $L$ [nm] | $\alpha$ [%] |
| --- | --- | --- | --- |
| Hopping molecules | $0.412 \pm 0.083$ | $1038 \pm 56$ | $31.4 \pm 0.8$ |
| Diffusing molecules | $0.046 \pm 0.002$ | $192 \pm 13$ | $68.6 \pm 0.4$ |

## Notes

### Competing Interest Statement

The authors have declared no competing interest.

## REFERENCES

Axelrod, D. (2001). Total internal reflection fluorescence microscopy in cell biology. Traffic Cph. Den. 2, 764–774.

Belanis, L., Plowman, S. J., Rotblat, B., Hancock, J. F. and Kloog, Y. (2008). Galectin-1 is a novel structural component and a major regulator of h-ras nanoclusters. Mol. Biol. Cell 19, 1404–1414.

Bello-Perez, M., Pereiro, P., Coll, J., Novoa, B., Perez, L. and Falco, A. (2020). Zebrafish C-reactive protein isoforms inhibit SVCV replication by blocking autophagy through interactions with cell membrane cholesterol. Sci. Rep. 10, 566.

Bernardello, M., Gora, R. J., Van Hage, P., Castro-Olvera, G., Gualda, E. J., Schaaf, M. J. M. and Loza-Alvarez, P. (2021). Analysis of intracellular protein dynamics in living zebrafish embryos using light-sheet fluorescence single-molecule microscopy. Biomed. Opt. Express 12, 6205– 6227.

Bobroff, N. (1986). Position measurement with a resolution and noise-limited instrument. Rev. Sci. Instrum. 57, 1152–1157.

Campagnola, G., Nepal, K., Schroder, B. W., Peersen, O. B. and Krapf, D. (2015). Superdiffusive motion of membrane-targeting C2 domains. Sci. Rep. 5, 17721.

Chen, J., Zhang, Z., Li, L., Chen, B.-C., Revyakin, A., Hajj, B., Legant, W., Dahan, M., Lionnet, T., Betzig, E., et al. (2014). Single-molecule dynamics of enhanceosome assembly in embryonic stem cells. Cell 156, 1274–1285.

Chudakov, D. M., Matz, M. V., Lukyanov, S. and Lukyanov, K. A. (2010). Fluorescent Proteins and Their Applications in Imaging Living Cells and Tissues. Physiol. Rev. 90, 1103–1163.

Dittrich, P. S. and Schwille, P. (2001). Photobleaching and stabilization of. fluorophores used for single-molecule analysis. with one- and two-photon excitation. Appl. Phys. B 73, 829–837.

Donnert, G., Eggeling, C. and Hell, S. W. (2007). Major signal increase in fluorescence microscopy through dark-state relaxation. Nat. Methods 4, 81– 86.

Donnert, G., Eggeling, C. and Hell, S. W. (2009). Triplet-relaxation microscopy with bunched pulsed excitation. Photochem. Photobiol. Sci. Off. J. Eur. Photochem. Assoc. Eur. Soc. Photobiol. 8, 481–485.

Eggeling, C., Widengren, J., Rigler, R. and Seidel, C. A. (1998). Photobleaching of Fluorescent Dyes under Conditions Used for Single-Molecule Detection: Evidence of Two-Step Photolysis. Anal. Chem. 70, 2651–2659.

Fujiwara, T., Ritchie, K., Murakoshi, H., Jacobson, K. and Kusumi, A. (2002). Phospholipids undergo hop diffusion in compartmentalized cell membrane. J. Cell Biol. 157, 1071–1081.

Garcia-Parajo, M. F., Cambi, A., Torreno-Pina, J. A., Thompson, N. and Jacobson, K. (2014). Nanoclustering as a dominant feature of plasma membrane organization. J. Cell Sci. 127, 4995–5005.

Gebhardt, J. C. M., Suter, D. M., Roy, R., Zhao, Z. W., Chapman, A. R., Basu, S., Maniatis, T. and Xie, X. S. (2013). Single-molecule imaging of transcription factor binding to DNA in live mammalian cells. Nat. Methods 10, 421–426.

Gora, R. J., de Jong, B., van Hage, P., Rhiemus, M. A., van Steenis, F., van Noort, J., Schmidt, T. and Schaaf, M. J. M. (2021). Analysis of the H-Ras mobility pattern in vivo shows cellular heterogeneity inside epidermal tissue. Dis. Model. Mech. dmm.049099.

Gore, A. V., Pillay, L. M., Galanternik, M. V. and Weinstein, B. M. (2018). The zebrafish: A fintastic model for hematopoietic development and disease. WIREs Dev. Biol. 7, e312.

Graham, D. J. L., Tseng, S.-F., Hsieh, J.-T., Chen, D. J. and Alexandrakis, G. (2015). Dependence of Two-Photon eGFP Bleaching on Femtosecond Pulse Spectral Amplitude and Phase. J. Fluoresc. 25, 1775–1785.

Groeneweg, F. L., Royen, M. E. van, Fenz, S., Keizer, V. I. P., Geverts, B., Prins, J., Kloet, E. R. de, Houtsmuller, A. B., Schmidt, T. S. and Schaaf, M. J. M. (2014). Quantitation of Glucocorticoid Receptor DNA-Binding Dynamics by Single-Molecule Microscopy and FRAP. PLOS ONE 9, e90532.

Ha, T., Ting, A. Y., Liang, J., Caldwell, W. B., Deniz, A. A., Chemla, D. S., Schultz, P. G. and Weiss, S. (1999). Single-molecule fluorescence spectroscopy of enzyme conformational dynamics and cleavage mechanism. Proc. Natl. Acad. Sci. U. S. A. 96, 893–898.

Hancock, J. F. and Parton, R. G. (2005). Ras plasma membrane signaling platforms. Biochem. J. 389, 1–11.

Harms, G. S., Cognet, L., Lommerse, P. H. M., Blab, G. A. and Schmidt, T. (2001). Autofluorescent Proteins in Single-Molecule Research: Applications to Live Cell Imaging Microscopy. Biophys. J. 80, 2396–2408.

Herrero, A., Matallanas, D. and Kolch, W. (2016). The spatiotemporal regulation of RAS signaling. Biochem. Soc. Trans. 44, 1517–1522.

Keizer, V. I. P., Coppola, S., Houtsmuller, A. B., Geverts, B., van Royen, M. E., Schmidt, T. and Schaaf, M. J. M. (2019). Repetitive switching between DNA-binding modes enables target finding by the glucocorticoid receptor. J. Cell Sci. 132, jcs217455.

Kugler, E. C., van Lessen, M., Daetwyler, S., Chhabria, K., Savage, A. M., Silva, V., Plant, K., MacDonald, R. B., Huisken, J., Wilkinson, R. N., et al. (2019). Cerebrovascular endothelial cells form transient Notch-dependent cystic structures in zebrafish. EMBO Rep. 20, e47047.

Kusumi, A., Sako, Y. and Yamamoto, M. (1993). Confined lateral diffusion of membrane receptors as studied by single particle tracking (nanovid microscopy). Effects of calcium-induced differentiation in cultured epithelial cells. Biophys. J. 65, 2021–2040.

Kusumi, A., Fujiwara, T. K., Chadda, R., Xie, M., Tsunoyama, T. A., Kalay, Z., Kasai, R. S. and Suzuki, K. G. N. (2012). Dynamic Organizing Principles of the Plasma Membrane that Regulate Signal Transduction: Commemorating the Fortieth Anniversary of Singer and Nicolson’s Fluid-Mosaic Model. Annu. Rev. Cell Dev. Biol. 28, 215–250.

Kwik, J., Boyle, S., Fooksman, D., Margolis, L., Sheetz, M. P. and Edidin, M. (2003). Membrane cholesterol, lateral mobility, and the phosphatidylinositol 4,5-bisphosphate-dependent organization of cell actin. Proc. Natl. Acad. Sci. U. S. A. 100, 13964–13969.

Lemmon, M. A. (2008). Membrane recognition by phospholipid-binding domains. Nat. Rev. Mol. Cell Biol. 9, 99–111.

Li, H. and Vaughan, J. C. (2018). Switchable Fluorophores for Single-Molecule Localization Microscopy. Chem. Rev. 118, 9412–9454.

Li, W., Han, M. and Guan, K. L. (2000). The leucine-rich repeat protein SUR-8 enhances MAP kinase activation and forms a complex with Ras and Raf. Genes Dev. 14, 895–900.

Lingwood, D. and Simons, K. (2010). Lipid Rafts As a Membrane-Organizing Principle. Science 327, 46–50.

Lommerse, P. H. M., Blab, G. A., Cognet, L., Harms, G. S., Snaar-Jagalska, B. E., Spaink, H. P. and Schmidt, T. (2004). Single-Molecule Imaging of the H-Ras Membrane-Anchor Reveals Domains in the Cytoplasmic Leaflet of the Cell Membrane. Biophys. J. 86, 609–616.

Lommerse, P. H. M., Snaar-Jagalska, B. E., Spaink, H. P. and Schmidt, T. (2005). Single-molecule diffusion measurements of H-Ras at the plasma membrane of live cells reveal microdomain localization upon activation. J. Cell Sci. 118, 1799–1809.

Lommerse, P. H. M., Vastenhoud, K., Pirinen, N. J., Magee, A. I., Spaink, H. P. and Schmidt, T. (2006). Single-Molecule Diffusion Reveals Similar Mobility for the Lck, H-Ras, and K-Ras Membrane Anchors. Biophys. J. 91, 1090–1097.

Lu, J.-W., Ho, Y.-J., Yang, Y.-J., Liao, H.-A., Ciou, S.-C., Lin, L.-I. and Ou, D.-L. (2015). Zebrafish as a disease model for studying human hepatocellular carcinoma. World J. Gastroenterol. 21, 12042–12058.

Luo, F., Qin, G., Xia, T. and Fang, X. (2020). Single-Molecule Imaging of Protein Interactions and Dynamics. Annu. Rev. Anal. Chem. Palo Alto Calif 13, 337–361.

Malumbres, M. and Barbacid, M. (2003). RAS oncogenes: the first 30 years. Nat. Rev. Cancer 3, 459–465.

Miller, H., Zhou, Z., Shepherd, J., Wollman, A. J. M. and Leake, M. C. (2018). Single-molecule techniques in biophysics: a review of the progress in methods and applications. Rep. Prog. Phys. Phys. Soc. G. B. 81, 024601.

Mione, M. C. and Trede, N. S. (2010). The zebrafish as a model for cancer. Dis. Model. Mech. 3, 517–523.

Morone, N., Nakada, C., Umemura, Y., Usukura, J. and Kusumi, A. (2008). Three-dimensional molecular architecture of the plasma-membrane-associated cytoskeleton as reconstructed by freeze-etch electron tomography. Methods Cell Biol. 88, 207–236.

Murakoshi, H., Iino, R., Kobayashi, T., Fujiwara, T., Ohshima, C., Yoshimura, A. and Kusumi, A. (2004). Single-molecule imaging analysis of Ras activation in living cells. Proc. Natl. Acad. Sci. 101, 7317–7322.

Murase, K., Fujiwara, T., Umemura, Y., Suzuki, K., Iino, R., Yamashita, H., Saito, M., Murakoshi, H., Ritchie, K. and Kusumi, A. (2004). Ultrafine membrane compartments for molecular diffusion as revealed by single molecule techniques. Biophys. J. 86, 4075–4093.

Nickels, J. D., Smith, J. C. and Cheng, X. (2015). Lateral organization, bilayer asymmetry, and inter-leaflet coupling of biological membranes. Chem. Phys. Lipids 192, 87–99.

Niesner, R., Andresen, V., Neumann, J., Spiecker, H. and Gunzer, M. (2007). The power of single and multibeam two-photon microscopy for high-resolution and high-speed deep tissue and intravital imaging. Biophys. J. 93, 2519–2529.

Patterson, G. H. and Piston, D. W. (2000). Photobleaching in two-photon excitation microscopy. Biophys. J. 78, 2159–2162.

Phillips, Z. F., Dean, S., Recht, B. and Waller, L. (2020). High-throughput fluorescence microscopy using multi-frame motion deblurring. Biomed. Opt. Express 11, 281–300.

Plowman, S. J., Muncke, C., Parton, R. G. and Hancock, J. F. (2005). H-ras, K-ras, and inner plasma membrane raft proteins operate in nanoclusters with differential dependence on the actin cytoskeleton. Proc. Natl. Acad. Sci. 102, 15500–15505.

Prior, I. A., Harding, A., Yan, J., Sluimer, J., Parton, R. G. and Hancock, J. F. (2001). GTP-dependent segregation of H-ras from lipid rafts is required for biological activity. Nat. Cell Biol. 3, 368–375.

Prior, I. A., Muncke, C., Parton, R. G. and Hancock, J. F. (2003). Direct visualization of Ras proteins in spatially distinct cell surface microdomains. J. Cell Biol. 160, 165–170.

Schaaf, M. J. M., Koopmans, W. J. A., Meckel, T., van Noort, J., Snaar-Jagalska, B. E., Schmidt, T. S. and Spaink, H. P. (2009). Single-Molecule Microscopy Reveals Membrane Microdomain Organization of Cells in a Living Vertebrate. Biophys. J. 97, 1206–1214.

Schütz, G. J., Schindler, H. and Schmidt, T. (1997a). Single-molecule microscopy on model membranes reveals anomalous diffusion. Biophys. J. 73, 1073–1080.

Schütz, G. J., Schindler, H. and Schmidt, T. (1997b). Single-molecule microscopy on model membranes reveals anomalous diffusion. Biophys. J. 73, 1073–1080.

Seefeldt, B., Kasper, R., Seidel, T., Tinnefeld, P., Dietz, K.-J., Heilemann, M. and Sauer, M. (2008). Fluorescent proteins for single-molecule fluorescence applications. J. Biophotonics 1, 74–82.

Semrau, S. and Schmidt, T. (2007). Particle Image Correlation Spectroscopy (PICS): Retrieving Nanometer-Scale Correlations from High-Density Single-Molecule Position Data. Biophys. J. 92, 613–621.

Shalom-Feuerstein, R., Plowman, S. J., Rotblat, B., Ariotti, N., Tian, T., Hancock, J. F. and Kloog, Y. (2008). K-ras nanoclustering is subverted by overexpression of the scaffold protein galectin-3. Cancer Res. 68, 6608–6616.

Shashkova, S. and Leake, M. C. (2017). Single-molecule fluorescence microscopy review: shedding new light on old problems. Biosci. Rep. 37, BSR20170031.

Silva, M. C. G. da, Silva, J. F. da, Santos, T. P., Silva, N. P. C. da, Santos, A. R. D., Andrade, A. L. C. de, Souza, E. H. L. da S., Sales Cadena, M. R., Sá, F. B. de, Silva Junior, V. A. da, et al. (2019). The complexation of steroid hormones into cyclodextrin alters the toxic effects on the biological parameters of zebrafish (Danio rerio). Chemosphere 214, 330–340.

Soeller, C. and Cannell, M. B. (1999). Two-photon microscopy: imaging in scattering samples and three-dimensionally resolved flash photolysis. Microsc. Res. Tech. 47, 182–195.

Suzuki, K., Ritchie, K., Kajikawa, E., Fujiwara, T. and Kusumi, A. (2005). Rapid Hop Diffusion of a G-Protein-Coupled Receptor in the Plasma Membrane as Revealed by Single-Molecule Techniques. Biophys. J. 88, 3659–3680.

Tinevez, J.-Y., Perry, N., Schindelin, J., Hoopes, G. M., Reynolds, G. D., Laplantine, E., Bednarek, S. Y., Shorte, S. L. and Eliceiri, K. W. (2017). TrackMate: An open and extensible platform for single-particle tracking. Methods San Diego Calif 115, 80–90.

Travers, T., Colin, V. G., Loumaigne, M., Barillé, R. and Gindre, D. (2020). Single-Particle Tracking with Scanning Non-Linear Microscopy. Nanomater. Basel Switz. 10, E1519.

Umemura, Y. M., Vrljic, M., Nishimura, S. Y., Fujiwara, T. K., Suzuki, K. G. N. and Kusumi, A. (2008). Both MHC class II and its GPI-anchored form undergo hop diffusion as observed by single-molecule tracking. Biophys. J. 95, 435–450.

van den Broek, B., Ashcroft, B., Oosterkamp, T. H. and van Noort, J. (2013). Parallel nanometric 3D tracking of intracellular gold nanorods using multifocal two-photon microscopy. Nano Lett. 13, 980–986.

Yasui, M., Matsuoka, S. and Ueda, M. (2014). PTEN Hopping on the Cell Membrane Is Regulated via a Positively-Charged C2 Domain. PLoS Comput. Biol. 10, e1003817.

Yokota, H. (2020). Fluorescence microscopy for visualizing single-molecule protein dynamics. Biochim. Biophys. Acta Gen. Subj. 1864, 129362.

Zhou, Y. and Hancock, J. F. (2015). Ras nanoclusters: Versatile lipid-based signaling platforms. Biochim. Biophys. Acta 1853, 841–849.

Zhou, Y. and Hancock, J. F. (2021). Super-Resolution Imaging and Spatial Analysis of RAS on Intact Plasma Membrane Sheets. Methods Mol. Biol. Clifton NJ 2262, 217–232.

Zhou, Y., Prakash, P., Gorfe, A. A. and Hancock, J. F. (2018). Ras and the Plasma Membrane: A Complicated Relationship. Cold Spring Harb. Perspect. Med. 8, a031831.

